# Membrane Proteins Have Distinct Fast Internal Motion and Residual Conformational Entropy

**DOI:** 10.1101/2020.04.02.022368

**Authors:** Evan S. O’Brien, Brian Fuglestad, Henry J. Lessen, Matthew A. Stetz, Danny W. Lin, Bryan S. Marques, Kushol Gupta, Karen G. Fleming, A. Joshua Wand

## Abstract

For a variety of reasons, the internal motions of integral membrane proteins have largely eluded comprehensive experiential characterization. Here, the fast side chain dynamics of the 7-transmembrane helix protein sensory rhodopsin II and the beta-barrel bacterial outer membrane channel protein W have been characterized in lipid bilayers and detergent micelles by solution NMR relaxation techniques. Though of quite different topologies, both proteins are found to have a similar and striking distribution of methyl-bearing amino acid side chain motion that is independent of membrane mimetic. The methyl-bearing side chains of both proteins, on average, are more dynamic in the ps-ns time regime than any soluble protein characterized to date. Approximately one third of methyl-bearing side chains exhibit extreme rotameric averaging on this timescale. Accordingly, both proteins retain an extraordinary residual conformational entropy in the folded state, which provides a counterbalance to the absence of the hydrophobic effect that normally stabilizes the folded state of water-soluble proteins. Furthermore, the large reservoir of conformational entropy that is observed provides the potential to greatly influence the thermodynamics underlying a plethora of membrane protein functions including ligand binding, allostery and signaling.

## Main Text

The motions of amino acid side chains of proteins are important for understanding the connection between energetics, structure and function in these complex macromolecules. For example, the conformational entropy manifested in subnanosecond motion can be a pivotal contribution to the thermodynamics of molecular recognition by proteins.^1^ Furthermore, the dynamical disorder of side chains in the sub-nanosecond time regime is heterogeneously distributed throughout the protein, which has significant implications for important aspects of their function.^2^ Three types or classes of motion have been discerned thus far by NMR relaxation approaches: highly restricted motion within a single rotameric well (termed the ω-class), larger excursions within a rotameric well that are accompanied by occasional rotameric interconversion (α-class); and motions involving more extensive interconversion between two rotameric states (J-class).^2^ These classes of motion are often resolved as distinct statistical distributions of NMR-derived generalized order parameters.^2,3^ Importantly, ligand binding can result in redistribution of side chain motions.^1,4,5^ A quantitative interpretation of the dynamical response provides insights into the thermodynamics underlying molecular recognition and indicates that the associated changes in conformational entropy often have an important role in the overall free energy of ligand binding.^1,6^ This view has been obtained entirely with soluble proteins.^1^ For a variety of reasons, membrane proteins have largely eluded similar investigations. Here we report the first comprehensive experimental characterization of the fast internal motions of two integral membrane proteins: sensory rhodopsin II (pSRII) and outer membrane protein W (OmpW). Both proteins display a similar distribution of fast side chain motion that is markedly different from that seen soluble proteins and corresponds to an unusually high residual conformational entropy.

Our findings are particularly remarkable because OmpW and pSRII represent the extremes of integral membrane protein topologies. pSRII is a 7-transmembrane helix homologue of G-protein coupled receptor proteins. The structure of isolated pSRII has been determined by crystallography (Fig. 1A)^7^ in lipid bilayers and by solution NMR spectroscopy in micelles^8^ and are in good agreement.^8^ In contrast, OmpW is a β-barrel membrane protein found in the outer membrane of *E. coli* and other Gram-negative bacteria. Like pSRII, the structure of OmpW has been solved using both x-ray crystallography and solution NMR spectroscopy.^9,10^ It contains 8 transmembrane β-strands that form a barrel structure in the membrane (Fig. 1B). Half of the inter-strand connections are short turns while the other four connections are large, loop structures. The functions of pSRII and OmpW are also distinct. pSRII mediates negative phototaxis in response to absorption of blue light. The 13-*trans*-*cis* isomerization of the attached retinal cofactor activates a two-component signaling system through its bound transducer partner protein.^11^ Though its precise function has not been confirmed, OmpW has been implicated in multiple cellular processes, such as transport of hydrophobic substrates,^9^ iron uptake,^12^ and antibiotic resistance.^13^ Given these structural and functional differences, the large reservoir of conformational entropy observed here in both points to a general role for fast motions in the thermodynamics governing membrane protein stability, folding and function.

**Fig. 1.**
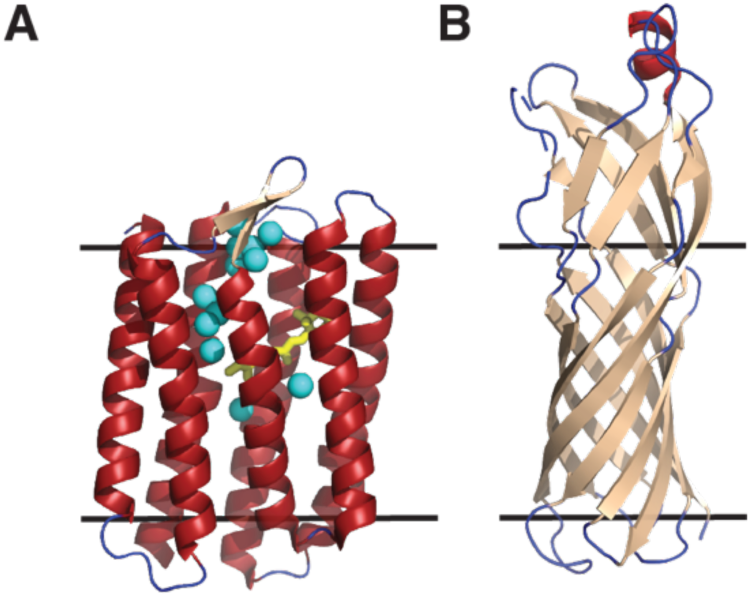
Structural folds of pSRII and OmpW. The crystal structures of pSRII (**A**) (PDB code 1H68) and OmpW (**B**) (PDB code 2F1T) are colored by secondary structural elements; α - helices in red, β - sheet in tan, and loops in blue. The black lines bracket approximate bilayer-spanning regions of the proteins. The retinal cofactor of pSRII is drawn as yellow sticks and structural water oxygen atoms are highlighted as cyan spheres.

## Results & Discussion

### Incorporation of integral membrane proteins into multiple membrane mimetics for NMR studies

We utilized a newly-developed growth and expression strategy (detailed elsewhere^14^) to generate fully ^15^N, ^13^CH_3_ ILVM labeled pSRII in the background of ∼75-80% deuteration. The labeling strategy involves expression in H_2_O and avoids “back exchange” of amide hydrogens that generally plagues preparation of membrane proteins generated by expression in D_2_O (Fig. S1A) while maintaining optimal methyl labeling (Fig. S1C) in a background of sufficient deuteration to permit quantitative measurement of methyl dynamics using cross correlated relaxation techniques (Fig. S1A, S1C).^15^ The presence of amide ^15^N-^1^H allows access to local backbone motion and determination of the molecular reorientation time of the protein.^16^ OmpW can be refolded in high yield and thus the traditional approach of expression during growth on bulk D_2_O media permitted a standard labeling strategy for ^15^N, ^13^CH_3_ ILV labeling in a background of carbon bonded deuterium (Fig. S1B & S1D). pSRII and OmpW were solubilized for NMR experiments in both detergent micelles (composed of c7-DHPC and SB3-12, respectively) as well as DMPC lipid bilayers in the form of q ∼ 1-1.1 bicelles where q is the molar ratio of long-chain DMPC and short chain DHPC. Small-angle X-ray scattering (SAXS) was used to confirm that a disc-like bicelle that maintained bilayer character was preserved in the presence of embedded protein.^17^ It has recently been shown that a feature of the scattering profile can be used to diagnose a transition from a disk-like bilayer containing bicelle to a mixed bicelle-micelle.^17^ For both pSRII and OmpW, addition of short-chain lipid results in a right-shift in a prominent peak feature in the mid-Q range (∼0.15 Å) as the bicelles transition into mixed micelles (Fig. S2).^17^ The onset of this transition begins below q ∼ 0.8 for both proteins. While this trend is clear for pSRII, the result for OmpW is rather muted, presumably due to the large extracellular domain that has serves to obscure the contribution of the bicelle to the scattering profile. Nevertheless, there is still a clear trend towards larger Q_max_ values for the main peak feature at lower bicelle q-ratio (Fig. S2B). In further support of the high-q bicelles used here, the Fas receptor transmembrane domain experiences a comparable transition upon modulation of bicelle q-value, as demonstrated by a series of detailed solution NMR characterizations of the protein itself (q ∼ 0.6-0.7).^18^ In summary, under the conditions used to obtain NMR relaxation data (q=1) both pSRII and OmpW are embedded in true bilayers.

Previous ^15^N relaxation studies of pSRII in DHPC micelles have indicated that the backbone of pSRII is essentially a rigid scaffold on the subnanosecond time scale.^19^ We repeated ^15^N longitudinal (R_1_) and transverse (R_2_) relaxation experiments to confirm a global reorientation time through a series of locally-resolved tumbling values, which is necessary for the accurate calculation of side chain dynamical parameters (21.9 ± 0.9 [micelle] and 28.9 ± 3.0 ns [bicelle]) (Fig. S3A & S3B). The backbone of pSRII is also largely silent in the slower µs-ms time regime as evidenced by R_1_·R_2_ products^20^ and generally flat ^15^N-dispersion profiles (Fig. S4A-D). Finally, Carr-Purcell-Meiboom-Gill (CPMG) methyl-dispersion experiments indicate that the methyl groups are devoid of µs – ms motions (not shown). Similarly, ^15^N backbone relaxation experiments conducted on OmpW in SB3-12 detergent micelles allowed determination of the global reorientation time (24.0 ± 0.9 ns [micelle] and 29.9 ± 2.2 ns [bicelle]) (Fig. S3C & S3D) and confirmed the lack of significant fast and slower timescale backbone motions (Fig. S4E & S4F).

### Membrane mimetic has little impact on fast internal motions of integral membrane proteins

Methyl cross-correlated relaxation experiments^15^ were then interpreted in the context of the Lipari-Szabo formalism,^21^ yielding squared generalized order parameter of the methyl group symmetry axis (O^2^_axis_) for nearly every resolved methyl resonance in pSRII and OmpW. O^2^_axis_ values range from one, corresponding to complete rigidity within the molecular frame, to zero, which effectively corresponds to isotropic disorder. Choice of lipid environment can be pivotal for interrogation of membrane protein structure, dynamics, & function.^22–26^ We have incorporated pSRII and OmpW into both detergent micelles as well as lipid bilayers in order to assess the potential role of membrane environment in membrane protein fast-timescale side chain dynamics. Due to their small size, micelles result in higher-quality, more comprehensive NMR data. Bilayers represent a more native-like environment for membrane proteins, yet form very large assemblies, limiting signal-to-noise. Comparisons of membrane protein dynamics between lipid conditions have been restricted to limited raw backbone relaxation data;^22,23,26^ we aimed here to quantitatively assess lipid environment effects by comparing dynamics *directly* through calculation of methyl order parameters. The NMR chemical shifts of pSRII & OmpW incorporated into micelles and bilayers are very comparable, though there are notable shifts in the OmpW ^13^C-methyl spectrum likely due to change in detergent/lipid head group (Fig. S1). Further, the NOESY patterns^8^ of pSRII prepared in DHPC micelles and DMPC bilayers in the form of bicelles are nearly identical, though maintenance of average parameters such as these are not necessarily indicative of conserved dynamics. To directly test this, we compared methyl O^2^_axis_ values for both proteins in the two environments. An excellent correlation of O^2^_axis_ values between the two lipid environments was observed for those methyl probes having data of suitable quality and that could be accurately mapped between membrane mimetics (R^2^ = 0.81 and 0.96 for pSRII and OmpW, respectively) (Fig. S5).

These comparisons suggest that fast-timescale motions are less sensitive to membrane mimetic than previously anticipated.^22,26^ Due to the apparent lack of dependence of fast-timescale dynamics on lipid environment for both membrane protein systems, we pursued more detailed investigations of micelle-incorporated proteins due to the improved data quality. Approximately 90/146 methyl probes in pSRII and 55/65 in OmpW had O^2^_axis_ values sufficiently determined with acceptable precision (error < 0.10) with an average error of ± 0.038 for pSRII (see Fig. S6A as well as the BMRB under accession number 27465) and ± 0.019 for OmpW (Fig. S6B)). Of note, the overwhelming majority of methyl groups not included for the analysis of pSRII are due to spectral overlap rather than low signal to noise. An “unbiased” analysis of all individual observed resonances in the pSRII methyl spectrum results in no notable deviations in either average O^2^_axis_ value or the resulting histogram (not shown). The O^2^_axis_ values reveal a surprising distribution of subnanosecond side chain motions in both membrane proteins.

### The extraordinary range of internal motion and entropy in integral membrane proteins

Both pSRII and OmpW display internal protein motion that is distinct from that previously observed for soluble proteins. The average O^2^_axis_ of pSRII is 0.36, which is unusually low and has not been observed previously in structured proteins (Fig. 2A). Ca^2+^-saturated calmodulin (CaM) is the next most dynamic protein characterized in this way (<O^2^_axis_> ∼ 0.43), largely due to a depopulated ω-band motional class.^4^ The cross-correlated relaxation experiments were best carried out at 50 °C for pSRII, which is somewhat higher than corresponding studies with the other proteins shown in Fig. 2A. Experiments at a more comparable but less optimal temperature of 35 °C gave an average change in O^2^_axis_ of +0.055, shifting the <O^2^_axis_ > at the lower temperature to 0.415. The apparent temperature dependence (dO^2^_axis_/dT) of pSRII is - 0.0037 K^-1^, which is slightly larger than that observed for a calmodulin complex^27^ and ubiquitin,^28^ and possibly hints at subtle differences in effective heat capacity between soluble and integral membrane proteins. Surprisingly, a comparable analysis of the topologically distinct OmpW demonstrates that it is *also* extraordinarily dynamic, with an average side chain methyl O^2^_axis_ value of 0.36 (Fig. 2A). Both membrane proteins measured in this manner are far more dynamic on average than any wild-type soluble protein investigated to date (Fig. 2A).

**Fig. 2.**
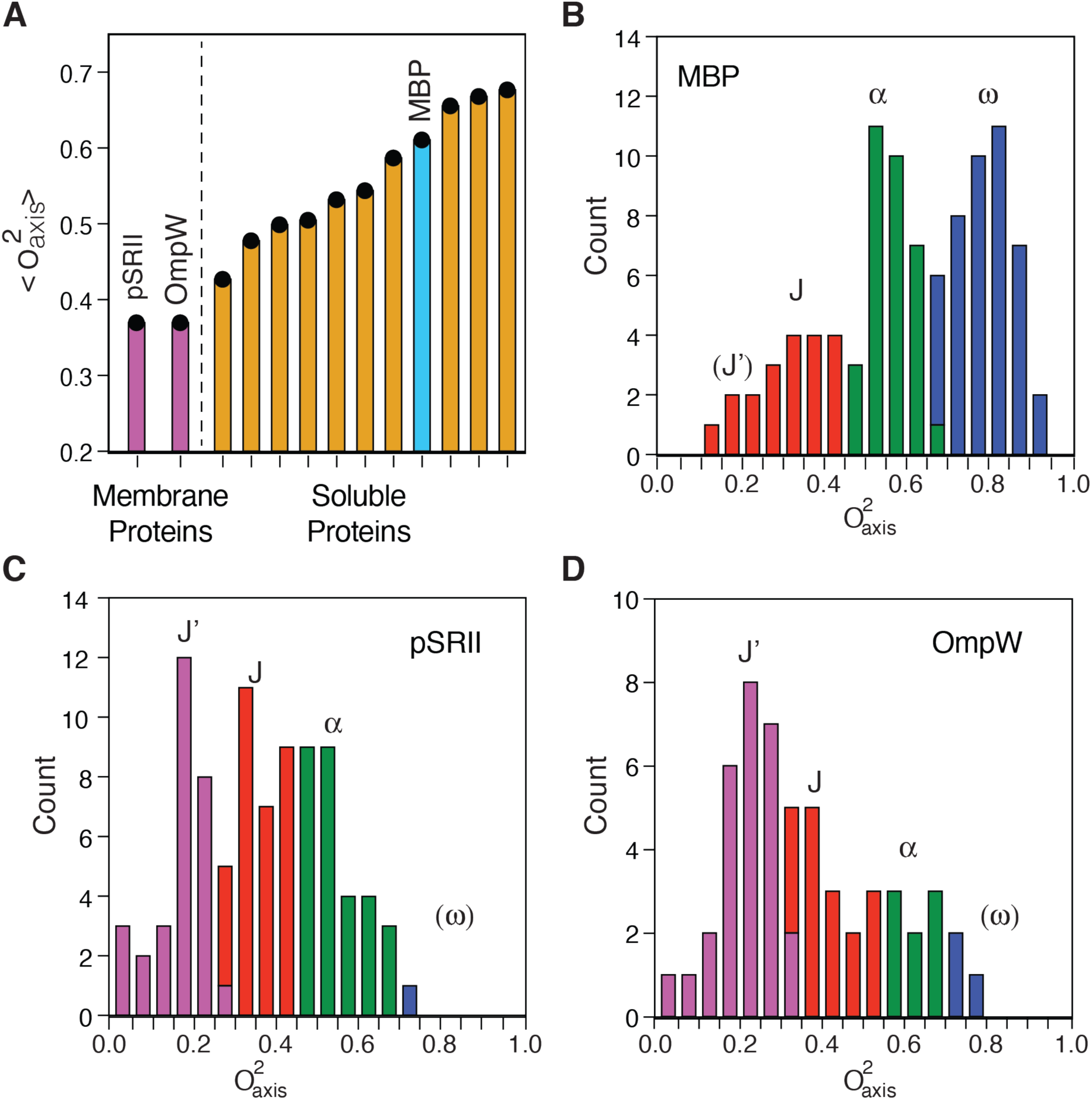
The distribution of fast side chain dynamics in membrane proteins is distinct from their soluble counterparts. (**A**) Average methyl-symmetry axis order parameters seen in soluble proteins and integral membrane proteins. Purple bars correspond to pSRII (50 °C) and OmpW (40 °C). The cyan bar corresponds to maltose binding protein (MPB), chosen to illustrate a representative distribution of soluble protein methyl-dynamics in (**B**). Orange bars correspond to other examples spanning the known range of soluble protein methyl-bearing side chain dynamics. The source data is described in Table S3. (**B**) Distribution of O^2^_axis_ values in MBP (Table S5). Shown to illustrate the distributions commonly seen in soluble proteins, particularly the segregation into three classes of underlying motion (J, red; α, green; ω, blue) as well as the emergence of a new class of motion in the two integral membrane proteins (J’, purple) seen in pSRII (**C**) and OmpW (**D**). Class boundaries were calculated using the k-means clustering algorithm. The rigid ω-class is effectively absent in both pSRII and OmpW. Class centers are indicated by the position of the class label. The newly observed J’-class is centered on an O^2^_axis_ of ∼0.21.

OmpW and pSRII also have a striking *distribution* of subnanosecond methyl-bearing side chain motion (Fig. 2C & 2D) that is distinct from that observed in soluble proteins (Fig. 2B). There is almost a complete absence of rigid methyl-bearing side chains (ω-class), which is generally significantly populated in soluble proteins (Fig. 2B). The populations of the J- and α-classes are roughly equivalent in the protein systems measured here. In addition, a previously undocumented and a qualitatively distinct class of motion is observed in these two membrane proteins. Approximately one third of the methyl bearing side chains exhibit an unusually high degree of dynamic disorder on the subnanosecond time scale. Analysis of the distribution of O^2^_axis_ values using a Bayesian statistical approach^3^ suggests two and three classes of motion with nearly the same likelihood. The three-class model seems more appropriate as two of the classes roughly match the centers of the J- and α-classes seen in soluble proteins; the centers of the J- and α-classes were further refined with a k-means clustering methodology and found to be 0.36 and 0.55, respectively, for pSRII. The J- and α-class centers for OmpW are 0.42 and 0.66. This roughly matches results for soluble proteins.^2^ Regardless, a new motional class (which we term J’) is centered at an O^2^_axis_ value of ∼0.21 for both proteins. Such a low order parameter suggests that two or more rotamers of each torsion angle are extensively sampled. Though molecular dynamics simulations do not reproduce the experimental O^2^_axis_ values quantitatively (Fig. S7), the highly dynamic J’ band of motions appears well reproduced by simulation in a bilayer. An analysis of those residues present in the J’ class in pSRII that are reasonably reproduced by simulation reinforces the notion that extensive rotameric interconversion of multiple torsion angles is required to achieve such low order parameters. This is in accord with theoretical considerations.^21,29^ The J’ class is also enriched relative to the overall average in methionine, leucine and isoleucine residues (73% versus 55%), which have two or more torsion angles to sample, and diminished in valine (27% versus 39%; only one non-terminal valine residue [Val101] is in the J’ class).

### Membrane proteins have a distinct statistical and structural distribution of side chain dynamics

We took advantage of previous methyl group assignments in pSRII to investigate the structural context of its side chain dynamics; this analysis reveals a number of interesting features. Methyl probes are depicted as spheres on the crystal structure^7^ and colored in a O^2^_axis_ gradient ranging from zero (red) to one (blue) in Fig. 3A. As in soluble protein systems, there is a heterogeneous distribution of disorder in O^2^_axis_ with no statistically significant (p-value > 0.2) spatial clustering of motional classes as revealed by the k-means approach,^30^ nor any significant correlation to any other obvious physical characteristics (Fig. S8). One might predict that methyl-bearing side chains exposed to lipid (or detergent) will tend to be more dynamic than residues more deeply buried in the structural core of the protein. However, pSRII does not display such “surface molten” behavior but rather has both highly dynamic and comparatively rigid probes exposed at the membrane (Fig. 3B) and aqueous surfaces of the protein (Fig. S8A). Methyl probes near what will serve as the interface with the transducer binding partner^31,32^ do not generally occupy the highly dynamic J’ motional class (<O^2^_axis_> = 0.45), though several relatively dynamic methyl probes sit at the extracellular side of the interface. This sort of “ordered interface” may serve to minimize entropy penalties upon interaction with the transducer. The region surrounding the retinal cofactor is not well sampled by well-determined methyl probes; many are sufficiently broadened by dipolar interactions with the ^1^H-rich retinal cofactor to degrade the quality of the relaxation measurements. However, Met109 Cε, which is ∼3.5 Å away and lies roughly perpendicular to the plane of conjugated double bonds, is one of the most rigid methyl groups in the protein (O^2^_axis_ = 0.68). The dynamic behavior, and changes therein, of side chains surrounding the retinal cofactor is likely to be important for efficient signal transduction. The unavailability of deuterated SB3-12 detergent thwarted collection of methyl assignment of sufficient quality to access resonance assignments and thus prevented a similar structural analysis of OmpW.

**Fig. 3.**
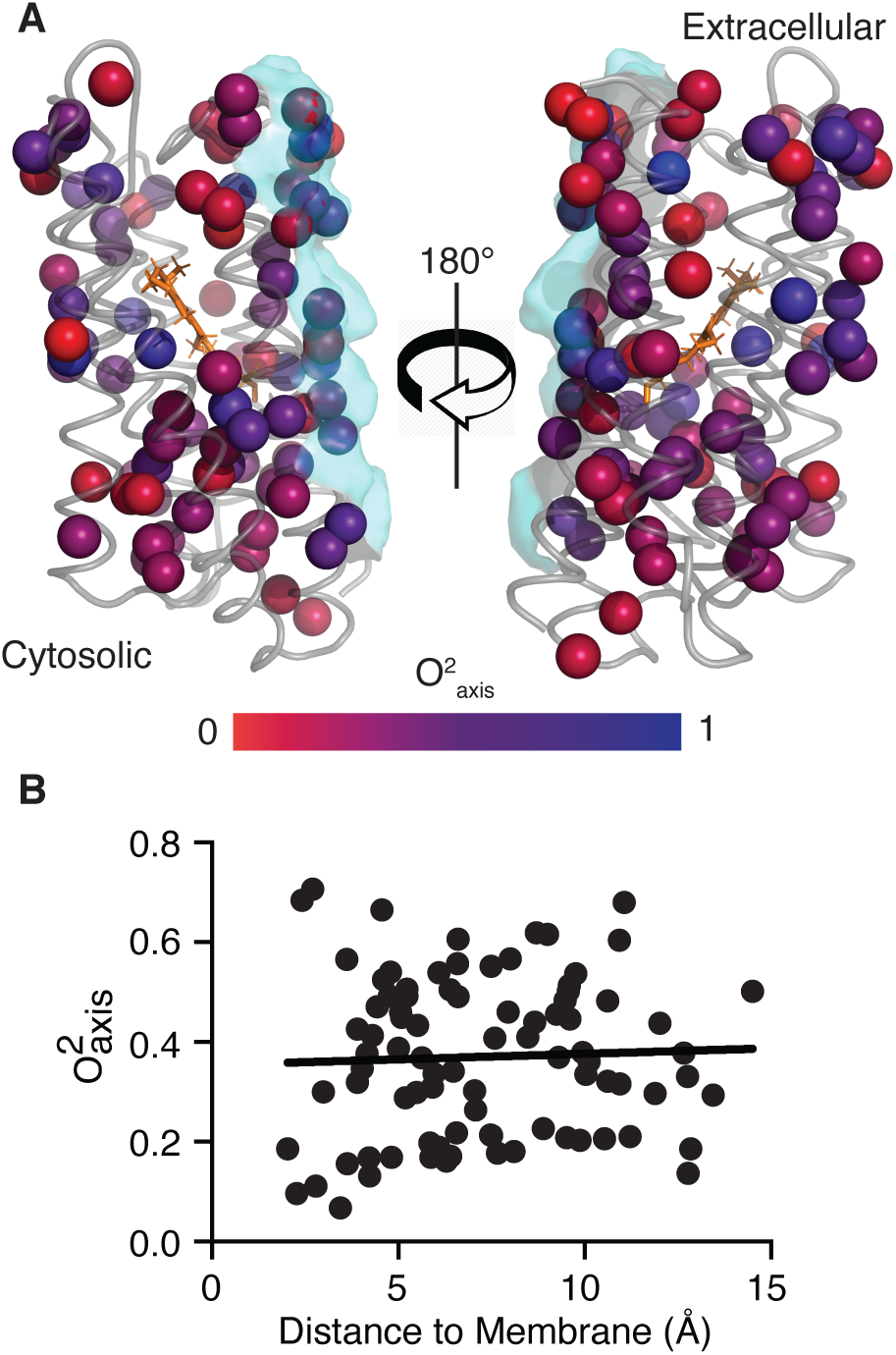
Spatial distribution of fast side chain motion in pSRII. (**A**) Shown is a ribbon representation of the crystal structure of pSRII (PDB code 1H68) on which spheres representing methyl probes are colored according to O^2^_axis_ value using a gradient of red for most dynamic (O^2^_axis_ = 0) to blue for most rigid (O^2^_axis_ = 1). The cytosolic and extracellular faces of the protein are noted. The retinal cofactor is shown as orange sticks and the surface of pSRII that forms the interface with its hTRII binding partner is shown as semi-transparent cyan. (**B**) Absence of a correlation of methyl dynamics with the distance to nearest lipid atom (R^2^ = 0.002).

The absence the ω-class of motions and emergence of the highly dynamic J’-class indicates that the methyl side chains are essentially fluid and raises a question of what interactions drive the formation of a stable tertiary structure. In this regard, it is interesting to note that small internal clusters of water molecules are commonly observed in crystal structures of GPCRs, which are structurally homologous to pSRII, and have been argued to play an important structural role as well as being actively involved in transitions between active/inactive states.^33^ In pSRII, crystallographic waters that are clustered between the extracellular β-sheet and the retinal and appear to support a buried arginine (Arg72) (Fig. 4A). This large water network lies near the extracellular face and is divided by a pair of hydrophobic residues (Ile197 and Val194) and Arg72, which is involved in hydrogen bonding interactions with both water clusters. Insight provided by methyl dynamics near this structural water network is limited to Val194 (O^2^_axis_ = 0.88 ± 0.11, not included in detailed statistics or Fig. 2 due to relatively high uncertainty) and Ile197 (O^2^_axis_ = 0.14), confirming the trend of heterogeneous methyl behavior throughout the protein tertiary structure. The presence of the water network in solution was confirmed using a methyl ^13^C-NOESY experiment. Strong negative NOE cross-peaks between the water resonance and methyl groups surrounding both deeply buried water clusters are seen (Fig. S9). The negative sign and strength of the NOEs indicate a relatively rigid protein-water interaction. Met15 which is proximal to the most buried water in cluster 2, Ile197 which lies between the water clusters, and Val68 which is near the extracellular-facing cluster 1 all display negative NOEs to the water resonance (Fig. S9). While many other methyl probes display NOEs to the water resonance, most are solvent exposed or potentially contaminated by nearby hydroxyl-containing residues, which may relay magnetization to water via hydrogen exchange.^34^ Unfortunately, the pSRII-containing micelle particle tumbles too slowly to permit measurement of the counterpart ROE that could potentially allow a more precise evaluation of the residence time of these buried waters.^35^ Notwithstanding the limitations of molecular dynamics to quantitatively reproduce experimentally determined order parameters in the current context, it is important to note that simulation suggests that nearly *all* buried polar side chains interacting with buried waters in both pSRII and OmpW are quite rigid on the fast-timescale, especially when compared to the dynamics of methyl bearing side chains in each protein (Fig. 4).

**Fig. 4.**
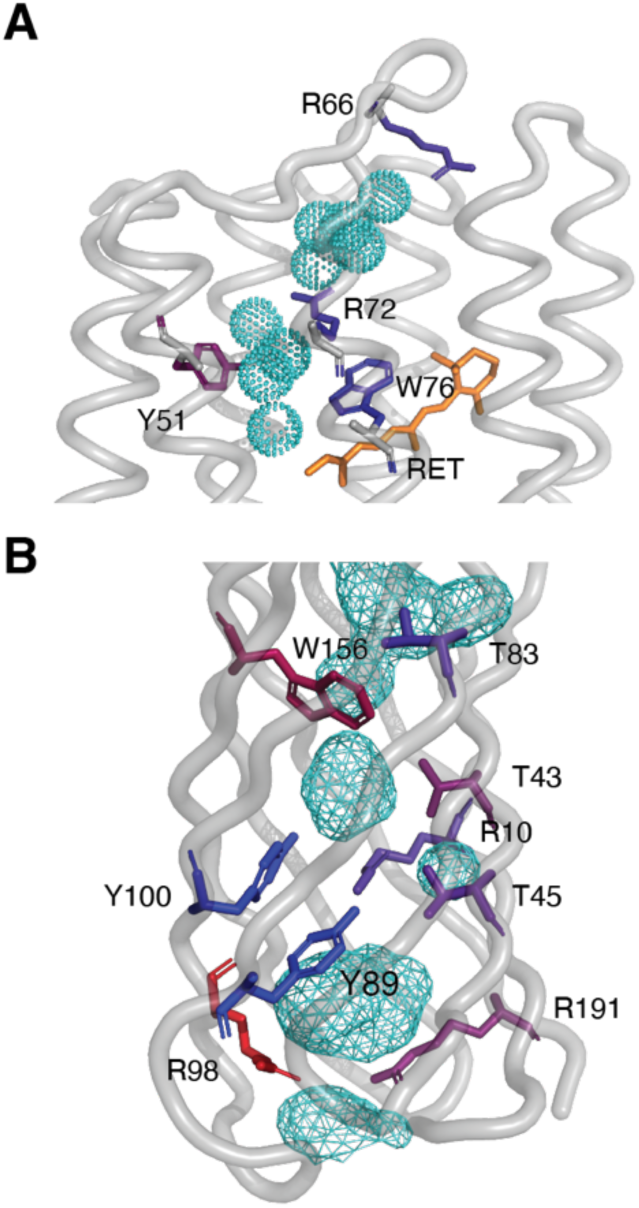
Polar cores of pSRII and OmpW. (**A**) Structural water oxygen atoms in the crystal structure (1H68) of pSRII are shown as cyan dotted spheres. Polar residues with side chains within hydrogen bonding distance of structural water molecules are depicted as sticks and are colored according to O^2^ values extracted from MD simulations according to the same red (O^2^ = 0.0) to blue (O^2^ = 1.0) scaling as Fig. 3. Order parameters were calculated via the Nε-Hε bond vector for arginine, The Cβ-Cγ2 vector for threonine, and the Cδ1-Hδ1 vector for tyrosine and tryptophan. The average polar side chain O^2^ in the core of pSRII is 0.70. (**B**) The crystal structure of OmpW (2F1T) is of insufficient resolution to resolve water molecules, so here the interior cavities are simply colored in cyan mesh. The average polar side chain O^2^ in the core of OmpW is 0.57.

## Discussion

In summary, the first comprehensive studies of fast side chain motion in integral membrane proteins presented here have revealed the existence of extensive sampling of microstates and point to a relatively high residual conformational entropy in pSRII and OmpW. pSRII and OmpW represent the topological extremes of integral membrane proteins and yet have the same dynamical signatures. As such they begin to suggest that their unusual dynamical character is a common feature of integral membrane proteins. Future work will address this question. Nevertheless, the high residual conformational entropy present in the native folded structures of these two proteins potentially impacts their stability, folding and function. For example, the stability of the folded state of an integral membrane protein,^36^ as for all proteins,^37^ results from a balance of forces. For soluble proteins, a dominant contribution to the stability of the folded state is the so-called hydrophobic effect that arises from the release of associated water from surfaces that are buried upon folding.^37^ Clearly, the high residual conformational entropy of the native states of both pSRII and OmpW contributes favorably to their stability. Furthermore, in a similar vein, most current models for folding of all alpha integral membrane proteins point to a two-step process where initial transfer of the unfolded polypeptide chain from water into the lipid bilayer is subsequently followed by folding to the final native state.^36,38^ Thus, in contrast to soluble proteins, folding of the polypeptide chain *within* the membrane lacks the general driving force of the hydrophobic effect i.e. the gain in solvent water entropy as the protein adopts a compact structure. The results presented here suggest that the extraordinary residual conformational entropy of the folded state helps avoid some of the penalty of organizing the tertiary fold within the membrane (Fig. 5). The large folding free energy of OmpW in large unilamellar vesicles (∼18 kcal mol^-1^) is also consistent with a minimal side chain entropy penalty upon adoption of the folded state.

**Fig. 5.**
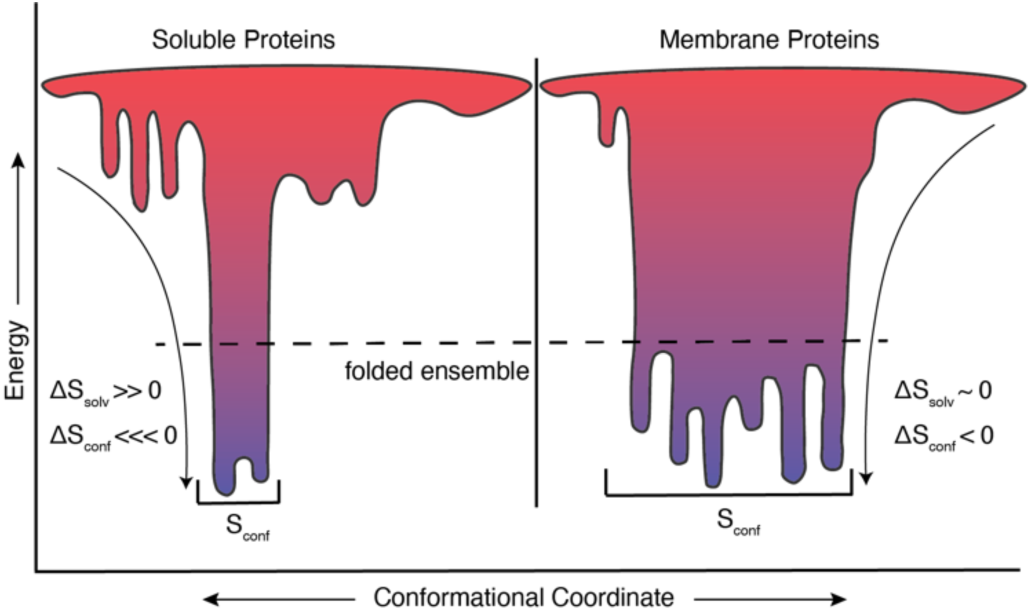
Soluble and membrane proteins utilize solvent and conformational entropies differently in the stabilization of their folded state ensembles. The folded state of soluble proteins is favored by a large increase in the entropy of water (ΔS_solv_, change in solvent entropy) and disfavored by a loss of chain and main and side chain entropy (ΔS_conf_, change in conformational conformational).^37^ In contrast, integral membrane proteins appear to mitigate the absence of a driving solvent water entropy through a diminished unfavorable contribution of reduced side chain conformational entropy.

The unusual dynamical character of pSRII also impacts our understanding of its mechanism of action and may serve as an informative counterpoint to G-protein coupled receptors (GPCRs) that are at the center of multicellular signaling. pSRII is a simple discrete binary switch and is apparently made so, in part, by a significant residual conformational entropy in the inactivated state that deepens the free energy well in which it resides. Conformational entropy manifested in methyl group motions, in concert with the water-mediated polar core interactions, simplifies its average structural character and would make a discrete transition upon activation possible. In contrast, non-olfactory GPCRs do not appear to occupy a localized structural ensemble and the lack of a single biophysical species has made detailed dynamical studies difficult.^39,40^ ^13^C methionine methyl labeling indicates that unliganded β_2_ adrenergic receptor (β_2_AR) displays extensive conformational exchange on the NMR chemical shift timescale (i.e. millisecond) indicating that barriers between microstates are large.^41^ GPCRs that are required to respond to an array of signaling events (i.e. binding of agonists, antagonists, G-proteins, β-arrestins) cycle between their various functional states. Indeed, a recent examination of A_2A_R reinforces this conclusion.^42^ Fast-timescale changes in structural water molecule positions near the retinal binding site in bacteriorhodopsin have been shown to facilitate these discrete transitions in functional state;^43^ highly dynamic methyl groups may be playing a similar role in activation. Finally, perturbations in these novel patterns of methyl dynamics in response to variations in membrane environment, binding partners, ligands, etc. are potentially functionally and thermodynamically important, and should yield further insights into the role of fast dynamics in membrane protein function and interaction.

## Acknowledgements

We thank K. G. Valentine and L. Liang for assistance throughout this project. We thank K. A. Sharp for helpful discussion.

## Funding

This work was supported by the G. Harold and Leila Y. Mathers Foundation (A.J.W) and R01 GM079440 (K.G.F.). E.S.O. and H.J.L are NIH predoctoral trainees (T32 GM008275 and T32 GM008403, respectively). This work used the Extreme Science and Engineering Discovery Environment (XSEDE) allocation MCB120050 (K.G.F.), which is supported by National Science Foundation grant number ACI-1548562 as well as computational resources from the Maryland Advanced Research Computing Center (MARCC) (K.G.F.).

## Author contributions

E.S.O. and A.J.W. designed the research. E.S.O., D.W.L., and H.J.L. refined protein purification methods. E.S.O., M.A.S. and H.J.L. performed all experiments; E.S.O and H.J.L. conducted and analyzed MD trajectories. B.F., H.J.L. and M.A.S. made and collected NMR experiments on bicelle samples. B.F. and K.G. collected SAXS experiments. E.S.O., M.A.S., H.J.L., K.G.F. and A.J.W. analyzed the data. E.S.O. and A.J.W. wrote the manuscript. A.J.W. and K.G.F. provided resources.

## Competing interests

The authors declare no competing financial interests.

## Data and materials availability

All data is available in the main text or the supplementary materials. Calculated O^2^_axis_ values for assigned methyl groups in pSRII with associated errors have been deposited to the Biological Magnetic Resonance Bank (BMRB) under accession number 27465.

## References

1. Caro, J. A. et al. Entropy in molecular recognition by proteins. Proc. Natl. Acad. Sci. U. S. A. 114, 6563–6568 (2017).

2. Igumenova, T. I., Frederick, K. K. & Wand, A. J. Characterization of the Fast Dynamics of Protein Amino Acid Side Chains Using NMR Relaxation in Solution. Chem. Rev. 106, 1672–1699 (2006).

3. Sharp, K. A., Kasinath, V. & Wand, A. J. Banding of NMR-derived methyl order parameters: Implications for protein dynamics. Proteins 82, 2106–17 (2014).

4. Lee, A. L., Kinnear, S. A. & Wand, A. J. Redistribution and loss of side chain entropy upon formation of a calmodulin-peptide complex. Nat. Struct. Biol. 7, 72–77 (2000).

5. Frederick, K. K., Marlow, M. S., Valentine, K. G. & Wand, A. J. Conformational entropy in molecular recognition by proteins. Nature 448, 325–329 (2007).

6. Wand, A. J. & Sharp, K. A. Measuring Entropy in Molecular Recognition by Proteins. Annu. Rev. Biophys. 47, 41–61 (2018).

7. Royant, A. et al. X-ray structure of sensory rhodopsin II at 2. 1-Å resolution. 98, 10131–10136 (2001).

8. Gautier, A., Mott, H. R., Bostock, M. J., Kirkpatrick, J. P. & Nietlispach, D. Structure determination of the seven-helix transmembrane receptor sensory rhodopsin II by solution NMR spectroscopy. Nat. Struct. Mol. Biol. 17, 768–774 (2010).

9. Hong, H., Patel, D. R., Tamm, L. K. & Van Den Berg, B. The outer membrane protein OmpW forms an eight-stranded β-barrel with a hydrophobic channel. J. Biol. Chem. 281, 7568–7577 (2006).

10. Horst, R., Stanczak, P. & Wuthrich, K. NMR Polypeptide Backbone Conformation of the E. coli Outer Membrane Protein W. Structure 22, 1204–1209 (2014).

11. Mironova, O. S. et al. Functional characterization of sensory rhodopsin II from Halobacterium salinarum expressed in Escherichia coli. FEBS Lett. 579, 3147–3151 (2005).

12. Catel-Ferreira, M. et al. The outer membrane porin OmpW of Acinetobacter baumannii is involved in iron uptake and colistin binding. FEBS Lett. 590, 224–231 (2016).

13. Lin, X., Yang, J., Peng, X. & Li, H. A Novel Negative Regulation Mechanism of Bacterial Outer Membrane Proteins in Response to Antibiotic Resistance research articles. J. Proteome Res. 9, 5952–5959 (2010).

14. O’Brien, E. S. et al. Improving yields of deuterated, methyl labeled protein by growing in H2O. J. Biomol. NMR 71, 263–273 (2018).

15. Sun, H., Kay, L. E. & Tugarinov, V. An optimized relaxation-based coherence transfer NMR experiment for the measurement of side-chain order in methyl-protonated, highly deuterated proteins. J. Phys. Chem. B 115, 14878–14884 (2011).

16. Jarymowycz, V. & Stone, M. J. Fast time scale dynamics of protein backbones: NMR relaxation methods, applications, and functional consequences. Chem. Rev. 106, 1624–71 (2006).

17. Caldwell, T. A. et al. Low-q Bicelles Are Mixed Micelles. J. Phys. Chem. Lett. 9, 4469–4473 (2018).

18. Piai, A., Fu, Q., Dev, J. & Chou, J. J. Optimal Bicelle q for Solution NMR Studies of Protein Transmembrane Partition. Chem. Eur. J. 23, 1361–1367 (2017).

19. Gautier, A., Kirkpatrick, J. P. & Nietlispach, D. Solution-state NMR spectroscopy of a seven-helix transmembrane protein receptor: Backbone assignment, secondary structure, and dynamics. Angew. Chemie - Int. Ed. 47, 7297–7300 (2008).

20. Kneller, J. M., Lu, M. & Bracken, C. An effective method for the discrimination of motional anisotropy and chemical exchange. J. Am. Chem. Soc. 124, 1852–1853 (2002).

21. Lipari, G. & Szabo, A. Model-free approach to the interpretation of nuclear magnetic resonance relaxation in macromolecules. 1. Theory and range of validity. J. Am. Chem. Soc. 104, 4546–4559 (1982).

22. Hagn, F., Etzkorn, M., Raschle, T. & Wagner, G. Optimized phospholipid bilayer nanodiscs facilitate high-resolution structure determination of membrane proteins. J. Am. Chem. Soc. 135, 1919–25 (2013).

23. Song, Y., Mittendorf, K. F., Lu, Z. & Sanders, C. R. Impact of Bilayer Lipid Composition on the Structure and Topology of the Transmembrane Amyloid Precursor C99 Protein. J. Am. Chem. Soc. 136, 4093–4096 (2014).

24. Frey, L., Lakomek, N.-A., Riek, R. & Bibow, S. Micelles, Bicelles, and Nanodiscs: Comparing the Impact of Membrane Mimetics on Membrane Protein Backbone Dynamics. Angew. Chemie Int. Ed. 56, 380–383 (2017).

25. Staus, D. P., Wingler, L. M., Pichugin, D., Prosser, R. S. & Lefkowitz, R. J. Detergent- and phospholipid-based reconstitution systems have differential effects on constitutive activity of G-protein–coupled receptors. J. Biol. Chem. 294, 13218–13223 (2019).

26. Bibow, S. & Hiller, S. A guide to quantifying membrane protein dynamics in lipids and other native-like environments by solution-state NMR spectroscopy. FEBS J. 286, 1610–1623 (2019).

27. Lee, A. L. & Wand, A. J. Microscopic origins of entropy, heat capacity and the glass transition in proteins. Nature 411, 501–504 (2001).

28. Song, X., Flynn, P. F., Sharp, K. A. & Wand, A. J. Temperature Dependence of Fast Dynamics in Proteins. Biophys. J. 92, L43–L45 (2007).

29. Wittebort, R. J. & Szabo, A. Theory of NMR relaxation in macromolecules : Restricted diffusion and jump models for multiple internal rotations in amino acid side chains Theory of NMR relaxation in macromolecules : Restricted diffusion and jump models for multiple internal rotations i. J. Chem. Phys. 69, 1722–1736 (1978).

30. Fu, Y. et al. Coupled motion in proteins revealed by pressure perturbation. J. Am. Chem. Soc. 134, 8543–8550 (2012).

31. Ishchenko, A. et al. New Insights on Signal Propagation by Sensory Rhodopsin II / Transducer Complex. Sci. Rep. 7, 41811 (2017).

32. Hippler-Mreyen, S. et al. Probing the sensory rhodopsin II binding domain of its cognate transducer by calorimetry and electrophysiology. J. Mol. Biol. 330, 1203–1213 (2003).

33. Yuan, S., Filipek, S., Palczewski, K. & Vogel, H. Activation of G-protein-coupled receptors correlates with the formation of a continuous internal water pathway. Nat. Commun. 5, 4733 (2014).

34. Otting, G. NMR studies of water bound to biological molecules. Prog. Nucl. Magn. Reson. Spectrosc. 31, 259–285 (1997).

35. Otting, G., Liepinsh, E. & Wüthrich, K. Protein hydration in aqueous solution. Science (80-.). 254, 974–980 (1991).

36. White, S. H. & Wimley, W. C. Membrane Protein Folding and Stability: Physical Principles. Annu. Rev. Biophys. Biomol. Struct. 28, 319–365 (1999).

37. Dill, K. A. Dominant Forces in Protein Folding. Biochemistry 29, 7133–7155 (1990).

38. Fleming, K. G. Energetics of Membrane Protein Folding. Annu. Rev. Biophys. 43, 233–55 (2014).

39. Tian, C. et al. Solution NMR Spectroscopy of the Human Vasopressin V2 Receptor, A G Protein-Coupled Receptor. J. Am. Chem. Soc. 127, 8010–8011 (2005).

40. Park, S. H. et al. Local and global dynamics of the G protein-coupled receptor CXCR1. Biochemistry 50, 2371–80 (2011).

41. Nygaard, R. et al. The dynamic process of β(2)-adrenergic receptor activation. Cell 152, 532–42 (2013).

42. Clark, L. D. et al. Ligand modulation of sidechain dynamics in a wild-type human GPCR. Elife 6, 1–27 (2017).

43. Nogly, P. et al. Retinal isomerization in bacteriorhodopsin captured by a femtosecond x-ray laser. Science (80-.). 361, 1–7 (2018).

