## Supplementary material for "Membrane Proteins Have Distinct Fast Internal Motion and Residual Conformational Entropy": Materials, Methods, and SI

2128 TAMU

College Station, TX 77843-2128

Telephone: (979)-845-2544 (Head Office)

**Materials and Methods**

**Sample Preparation**

*Expression of labeled pSRII* – A pET11a vector containing the gene for sensory rhodopsin II from *Natronomonas pharaonis* with a C-terminal histidine tag was freshly transformed into BL21(DE3) competent *E. coli* cells. An overnight culture was used to inoculate 1 L of 1.5x M9 medium in H_2_O with ^15^N labeling at an initial OD_600_ of ~0.1 at 37°C, pH 7.4, and 40% dissolved oxygen concentration. This first “phase” of growth contained 1 g ^15^NH_4_Cl, 2 g glucose, 200 mg yeast extract, 0.24 g MgSO_4_, 15 mg CaCl_2_, 15 mg ZnCl_2_, 15 mg FeSO_4_ and appropriate antibiotic. Temperature, pH, and oxygen were maintained or monitored in a New Brunswick BioFlo 115 cell culture system with a 3 L vessel. Cells were grown to saturation (OD_600_ ~ 2.4) as judged by the spike in dissolved oxygen concentration, at which point the temperature was dropped to 18°C, the pH was raised to 7.4, and 1 mL of 10 mM all-trans retinal in EtOH added along with isotopically labeled nutrients 10 minutes before induction of protein expression with 1 mM IPTG. The isotopically labeled nutrients included 1 g of ^2^H-D-glucose [Sigma-Aldrich], 1 g of ^15^N/^2^H algal-derived amino acids [Martek] depleted of isoleucine, leucine, valine, methionine, and tyrosine, 60 mg alpha-ketobutyrate [Sigma-Aldrich] and 100 mg alpha-ketoisovalerate [Cambridge Isotope Laboratories] both with ^13^CH_3_ methyl labeling of one isotopomer and ^12^CD labeling elsewhere, 50 mg phenyl-deuterated L-tyrosine [Sigma Aldrich], 50 mg ^13^CH_3_/^12^CD L-methionine [Cambridge Isotope Laboratories], 0.24 g MgSO_4_ and 15 mg CaCl_2_ were added. The algal-derived amino acids were depleted of I, L, V and M (and Y) amino acids by reverse phase HPLC; appropriately isotopically labeled precursors for ILV were then used along with labeled methionine and tyrosine as described above. Protein was expressed for ~3 hours at 18°C before glucose was fully depleted, with 1 mL of 10 mM all-trans retinal added every hour (30 µM total final concentration). Cells were spun down and frozen. Details of this expression strategy are described elsewhere.^14^ This quantity of ^2^H/^15^N amino acids results in ~80% final ^2^H incorporation.

*Purification of pSRII* - pSRII was purified in a slightly different manner to that described previously.^8,19^ Cell pellets (1 L culture) were resuspended in 20 mL 100 mM Tris-HCl pH 7.4, 400 mM NaCl and 20% glycerol. Lysozyme (20 mg), DNAse (4 mg), 800 μL 1 M MgCl_2_, 0.6 g dodecyl-maltopyranoside (DDM), and 2 EDTA-free protease inhibitor tablets [COEDTAF-RO Roche] were then added. The mixture was diluted to 40 mL with deionized H_2_O and rotary mixed in the dark overnight. The lysate was then centrifuged at 32,000 g for 45 minutes at 4°C. The insoluble pellet was discarded and 1 M imidazole was added to the supernatant to a final concentration of 20 mM (final volume ~ 40 mL) which was loaded at 0.7 mL/min onto a nickel column pre-equilibrated with 50 mM phosphate buffer pH 7.4, 150 mM NaCl, 10% glycerol, 20 mM imidazole, and 0.1% DDM detergent. The column was washed with 5 column volumes of the same buffer followed by five column volumes of the buffer supplemented with 750 mM NaCl and 50 mM imidazole. DDM detergent was exchanged for deuterated d_26_-diheptanoyl-phosphocholine (DHPC) detergent [CortecNet] as previously described^8^ and then eluted with 0.28% d_26_-DHPC in pH 6.0 50 mM phosphate, 300 mM imidazole with 50 mM NaCl and 0.02% sodium azide. The protein was exchanged into the same phosphate buffer without imidazole and concentrated to a final protein concentration not exceeding 500 µM.

*Expression of Labelled OmpW –* OmpW cloning and the plasmid construct used in this study has been previously described.^44^ Expression of isotopically labeled OmpW was carried out in an HMS *E coli* cell line using the *lac* expression system. A glycerol stock, kept at -80°C, was used to inoculate 3 mL starter growths. These starter cultures were grown in LB media supplemented with 0.1 mg/mL ampicillin at 37°C for 8 to 10 hrs. Cells were pelleted and resuspended in 1 mL of M9 media supplemented with 0.1 mg/mL ampicillin. Isotope incorporation for NMR studies was accomplished using ^15^NH_4_Cl (Cambridge Isotopes: NLM-467), uniform ^2^H or ^13^C/^2^H glucose (Cambridge Isotopes: DLM-2062, CDLM-3813), and D_2_O (Cambridge Isotopes: DLM-4). Selective methyl labeling was accomplished using standard ILV precursors.^45^ Labeling for protein used in methyl relaxation experiments utilized α-ketobutyrate and α-ketoisovalerate that lead to ^13^CH_3_ methyl labeling of one isotopomer and ^12^CD labeling elsewhere (Sigma-Aldrich 589276, Cambridge Isotopes CDLM 7317). Selective side chain labeling for methyl assignment experiments was accomplished using α-ketobutyrate and α-ketoisovalerate that lead to ^13^CH_3_ methyl labeling of one isotopomer and ^13^CD labeling elsewhere (Cambridge Isotopes: CDLM-4611, CDLM-8100). Two separate 25 mL growths were inoculated using 500 μL of resuspended cell solution. The 25 mL cultures were grown for ~17 hrs (to an OD between 0.8 and 0.9) and then diluted to 500 mL in M9 media. The 500 mL cultures were grown to an OD of ~0.6 (taking approximately 13 hours). At this point, the ILV precursors were added to the culture. After 1 hour, cells were induced by adding 1 mM IPTG. The induced culture was pelleted after ten hours and then frozen at -20°C.

*Purification of OmpW* – The procedure for inclusion body purification has been previously detailed for outer membrane proteins.^44,46^ It will be summarized briefly. The frozen cell pellet was thawed and resuspended in a lysis buffer (100 mM Tris-HCl, pH 8.0, 10 mM EDTA). The cells were then lysed using an Emulsiflex Cell Homogenizer. Following lysis, 87 μL of Brij-35 detergent was added to 25 mL of lysed cells. The lysed cell solution was then centrifuged at ~4,500 G for 20 minutes. The supernatant was discarded and the pellet resuspended in a wash buffer (100 mM Tris-HCl, pH 8.0, 1 mM EDTA) and centrifuged for an additional 20 minutes. The wash process was repeated, and then the inclusion body pellet was frozen at -20°C. The frozen inclusion body pellet was thawed and resuspended in 8 M urea buffered to pH 8.0 with 20 mM Tris-HCl. Final denatured OmpW concentration was 50 μM. The denatured OmpW was diluted dropwise into refolding buffer at 45°C being stirred at 600 RPM on a desktop stir plate. Final conditions for the refolding reaction were 4 μM OmpW, 20 mM Tris-HCl pH 8.0, and 6.5 mM SB3-12 detergent. After 12 hours, the reaction was quenched by adding acetate buffer to a final concentration of 40 mM. The system continued to stir for 1 hour and was then filtered using a 0.45-micron syringe filter. The filtered solution was then concentrated to a final volume of ~400 μL using 30 kD spin concentrators (EMD Millipore). Using the same concentrators, the buffer was exchanged into final conditions. Final buffer conditions were 40 mM phosphate buffer, pH 7.0, 6.5 mM SB3-12 detergent, and 10% (volume ratio) D_2_O. Final sample volume was 370 μL and concentration was assessed by absorbance at 280 nm and calculated using a molar extinction coefficient of 39,420 M^-1^. NMR sample was contained in a Shigemi tube (Sigma-Aldrich). Final protein concentrations for NMR samples were 275 μM (relaxation) and 670 μM (assignment).

*pSRII & OmpW Bicelle Incorporation* – Rather than exchange into d_26_-c7-DHPC detergent, for bicelles pSRII was similarly exchanged on-resin for 20 mM 1,2-dihexanoyl-sn-glycero-3-phosphocholine (c_6_-DHPC). pSRII in deuterated c6-DHPC micelles was eluted with buffer containing additional (20 mM) c6-DHPC and 300 mM imidazole. Eluted protein was buffer exchanged into the pSRII NMR buffer to remove imidazole. ^1^H-NMR was used to confirm the lack of detectable residual DDM. The long-chain lipid was prepared for bicelle incorporation by dissolving deuterated d_54_-DMPC to 50 mg/mL in chloroform. The amount needed to construct a sample of 1:1 DMPC:c6-DHPC (q=1.0) was added to a glass vial. A nitrogen stream was used to dry the DMPC into a thin layer against the glass. pSRII in c6-DHPC micelles was added to the dried lipid and vortexed. Five freeze-thaw cycles were performed by moving the sample between ice and a 42°C water bath in 5 minute intervals to further clarify the sample. NMR and SAXS experiments were performed as outlined below.

OmpW-containing bicelles were generated by resuspending 1,2-dimyristoyl-sn-glycero-3-phosphocholine (DMPC) in SB3-12 solubilized OmpW in pH 10, 40 mM borate buffer. DMPC was resuspended by performing 5 cycles of vortexting, heating to 42°C for 3 minutes, and chilling on ice for 3 minutes. Final concentrations of SB3-12 and DMPC were 23 mM and 27 mM, respectively, for a final bicelle q-value of ~1.1. ^15^N-TROSY spectra used in Fig. S1 were collected for OmpW in SB3-12 micelles at pH 10 with 40 mM borate buffer for comparison with bicelle-incorporated OmpW in order to have identical buffer conditions.

*MBP expression & purification* – Deuterated, ^15^N-labeled maltose binding protein, with ^13^CH_3_ labeled isoleucine, leucine, valine, and methionine residues was grown and purified as previously described.^47^ The same isotopic labeling strategy and scheme as for OmpW above was used for MBP.

**Small-angle X-ray Scattering**

pSRII and OmpW embedded in bicelles with q = 1.0 were used to construct a series of varying q ratios. The short-chain proportion was increased by adding c_6_-DHPC in the appropriate amount. The following ratios were measured via SAXS: q = 1.0, 0.8, 0.6, 0.4, 0.2. SAXS measurements were conducted using the 16ID-LiX Beamline at the National Synchrotron Light Source II, located at the Brookhaven National Laboratory (Upton, NY).^48^ Data were collected with the standard flow-cell-based solution scattering setup and at a wavelength of 1.0 Å in a three-camera configuration, yielding accessible scattering angle where 0.005 < q < 3.0 Å-1, where q is the momentum transfer, defined as q = 4π sin(θ)/ λ, where λ is the X-ray wavelength and 2θ is the scattering angle.. Radial averaging and *q*-conversion of data were performed using beamline-specific software, merging data from all three detectors used in the measurements. Transmission correction and background subtraction were performed to minimize the intensity of the hydrogen bond from water at ∼2.0 Å^−1^. 45 μL of sample were exposed for 5 replicates of 1-s each and averaged. Scattering curves were plotted using to determine if the expected shift in Q_max_ corresponded to bicellar shape in the q =1.0 samples.^17^

**NMR Spectroscopy**

*Micelle-incorporated pSRII* – All experiments were collected at 50 °C on Bruker AVANCE III spectrometers equipped with TXI cryoprobes. Most experiments were carried out at 750 MHz (^1^H). Assignments for the backbone amide and side chain methyl groups (methionine Cε, leucine Cδ1 and Cδ2, valine C𝛾1 and C𝛾2, and isoleucine Cδ1) were mapped from previous studies,^8,19^ and confirmed with a 3D NOESY ^1^H-^15^N TROSY experiment (backbone amide) and a 3D NOESY ^1^H-^13^C HMQC experiment (side chain methyl). These experiments were collected with non-uniform sampling at a sampling density of 15% with a Poisson-gap distribution.^49^ Correlation with previous assignments^8,19^ are excellent (R^2^ > 0.996). Two-dimensional ^13^C-HMQC and ^15^N-TROSY experiments were collected with 200 and 128 complex indirect points, respectively. A 3D NOESY – ^13^C,^1^H – HSQC experiment with 20% sampling density with a Poisson gap distribution and 200 ms mixing time was collected to observe the presence of specific methyl-water interactions in the core of the protein. All non-uniformly sampled data were reconstructed with iterative soft thresholding (istHMS)^49^ and processed in nmrPipe.^50^

^15^N-TROSY sampled ^15^N T_1_ relaxation experiments^51^ for the micelle-incorporated pSRII were collected at 750 MHz with relaxation delays of 0.000, 0.320 (2x), 0.560, 0.880, 1.200 (2x), and 1.600 and at 600 MHz with relaxation delays of 0.000, 0.320 (2x), 0.640, 0.960, 1.440 (2x), and 2.000 s. ^15^N-TROSY sampled T_1ρ_ relaxation experiments^51^ were collected at 750 MHz with relaxation delays of 0.001, 0.0077 (2x), 0.0141, 0.0218, 0.0304 (2x), and 0.0400 s at 600 MHz with relaxation delays of 0.001, 0.0067 (2x), 0.0124, 0.0190, 0.0266 (2x), and 0.035 s. Decay curves for resolved peaks in the ^15^N-TROSY spectrum were fitted to single-exponential decays with errors determined by deviations between duplicate delay values. T_1ρ_ values were converted to their corresponding T_2_ values using the ^15^N spin-lock RF field strength (ω_1_) and the individual offset from the ^15^N carrier in Hz (Ω) for each peak using Eq. 1 where tanθ = ω_1_/Ω.^51^

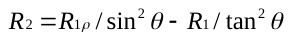
**(1)**

The obtained T_1_ and T_2_ values were used to determine the effective tumbling times for each residue as calculated:^52^

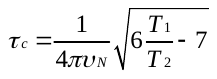
**(2)**

After excluding separately tumbling outliers from the distribution (see Fig. S3), the average was selected as the global reorientation time yielding values of 22.8 and 21.0 ns for data obtained at 600 and 750 MHz (^1^H), respectively. An average tumbling time of 21.9 ± 0.9 ns was used for analysis of cross-correlated relaxation data, which is in very good agreement with that obtained previously (21.3 ns)^19^ was used for analysis of the cross-correlated relaxation data. As further validation of the global tumbling time, a separately grown, purified, and prepared sample was analyzed in a similar manner, yielding values of 22.7 ns (600 MHz) and 21.3 ns (750 MHz).

Backbone amide^53^ and side chain methyl^54^ Carr-Purcell-Meiboom-Gill (CPMG) experiments were collected on micelle-incorporated pSRII at 750 MHz with various spacing between adjacent 180° pulses (1.00, 1.25, 1.67, 2.50, 3.33 (2x), 5.00, 6.67, 10.00, 20.00 ms intervals in a 40 ms constant time period for backbone amide CPMG and 1.00, 1.25, 1.67 (2x), 2.00, 2.50, 2.86, 3.33, 4.00, 5.00, 6.66, 10.00, and 20.00 ms spacings in a 20 ms constant time period for side chain methyl CPMG experiments) to detect the presence of intermediate timescale (µs – ms) motions.

Side chain methyl (ILVM) order parameter (O^2^_axis_) values were determined by cross-correlated relaxation experiments.^15^ Single- and triple-quantum ^1^H-^13^C coherence transfer experiments were collected at a series of delay times: 0.003, 0.005, 0.008 (2x), 0.012, 0.017, 0.022 (2x), 0.028, and 0.036 s. More scans (144/FID) for acquired for the triple quantum experiment than for the single-quantum experiment (64/FID) and the raw peak intensities normalized accordingly. Ratios of normalized peak intensities were fitted for η and δ values using Eq. 3 where *T* is the relaxation time:^15^

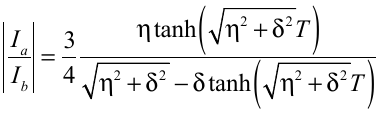
 **(3)**

O^2^_axis_ values were then determined for each methyl probe as in Eq. 4, using their fitted η values and the separately determined macromolecular reorientation time (𝜏_c_) of 21.9 ± 0.9 ns. For slow isotropic macromolecular tumbling:^15^

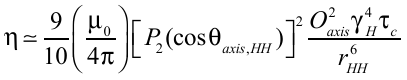
 **(4)**

where μ_0_ is the vacuum permittivity constant, γ_H_ is the proton gyromagnetic ratio, r_HH_ is the distance between protons within a methyl group (1.813 Å), θ_axis,HH_ is the angle between the methyl symmetry axis and a vector between a pair of methyl protons (90°), and P_2_(x) = ½(3x^2^ – 1). Uncertainties in the final O^2^_axis_ values were determined by 500 Monte Carlo simulations incorporating errors in fitted η values and tumbling time. Fitted η and δ values, χ^2^, local reduced χ^2^, final O^2^_axis_ values and their errors are provided in Table S1, and order parameter values and associated errors are deposited to the BMRB under accession number 27465. A comprehensive set of L-S order parameters were obtained from cross-correlated relaxation experiments at 50 °C. To examine the temperature sensitivity of motion, cross-correlated relaxation experiments were also carried out at 35 °C for the micelle-incorporated receptor. Because of an increased tumbling time (𝜏_c_ of 30.2 ± 0.9) at the lower temperature the sensitivity was greatly reduced and only a subset of peaks (~35) in the full spectrum could be analyzed quantitatively for comparison. A Bayesian statistical model^3^ where the data are analyzed for the presence of an unknown number (1-5) of motional classes was used to evaluate the distribution of O^2^_axis_ values. The 3-banded distribution was further explored with a k-means clustering algorithm described previously.^55^

*Bicelle-incorporated pSRII* – ^15^N-TROSY sampled ^15^N T_1_ relaxation experiments^51^ for the bicelle-incorporated pSRII were collected at 800 MHz with relaxation delays of 0.000, 0.160 (2x), 0.480, 1.280, 1.840 (2x), and 2.400. ^15^N-TROSY sampled T_1ρ_ relaxation experiments^51^ were collected at 800 MHz with relaxation delays of 0.001, 0.002 (2x), 0.006, 0.012, 0.02 (2x), and 0.0300 s. Decay curves for resolved peaks in the ^15^N-TROSY spectrum were fitted to single-exponential decays with errors determined by deviations between duplicate delay values. T_1ρ_ values were converted to their corresponding T_2_ values as above. Tumbling time was calculated as above, with the error being the standard deviation of the individual site tumbling time distribution.

Single- and triple-quantum ^1^H-^13^C coherence transfer experiments were collected at a series of delay times: 0.002, 0.003, 0.005 (2x), 0.007, 0.010, 0.013, 0.016 (2x), 0.019, and 0.022 s. More scans (144/FID) for acquired for the triple quantum experiment than for the single-quantum experiment (32/FID) and the raw peak intensities normalized accordingly. Side chain methyl order parameter values were calculated as above for the micelle-incorporated pSRII.

*Micelle-incorporated OmpW* – ^15^N-TROSY sampled ^15^N T_1_ relaxation experiments^51^ were collected at 600 MHz with relaxation delays of 0.000, 0.320 (2x), 0.640, 0.960, 1.440 (2x), 1.920, and 2.400 s. ^15^N-TROSY sampled T_1ρ_ relaxation experiments^51^ were also collected at 600 MHz with relaxation delays of 0.001, 0.0067 (2x), 0.0124, 0.0190, 0.0266 (2x), and 0.0350 s Decay curves for resolved peaks in the ^15^N-TROSY spectrum analyzed identically. Single- and triple-quantum ^1^H-^13^C coherence transfer experiments were collected at a series of delay times: 0.001, 0.003, 0.005 (2x), 0.007, 0.010, 0.013, 0.016, 0.019 (2x), 0.022, and 0.026 s. More scans (96/FID) for acquired for the triple quantum experiment than for the single-quantum experiment (32/FID) and the raw peak intensities normalized accordingly. Ratios of normalized peak intensities were fitted for η and δ values as described for pSRII above.

*Bicelle-incorporated OmpW* – ^15^N-TROSY sampled ^15^N T_1_ relaxation experiments^51^ were collected at 800 MHz with relaxation delays of 0.000, 0.160 (2x), 0.480, 1.280, 1.840 (2x), and 2.400 s. ^15^N-TROSY sampled T_1ρ_ relaxation experiments^51^ were also collected at 800 MHz with relaxation delays of 0.001, 0.002 (2x), 0.004, 0.008, 0.012, 0.016 (2x), 0.020, and 0.025 s. Decay curves for resolved peaks in the ^15^N-TROSY spectrum analyzed identically. Single- and triple-quantum ^1^H-^13^C coherence transfer experiments were collected at a series of delay times: 0.002, 0.003, 0.005 (2x), 0.007, 0.010, 0.013, 0.016 (2x), 0.019, and 0.022 s. More scans (96/FID) for acquired for the triple quantum experiment than for the single-quantum experiment (32/FID) and the raw peak intensities normalized accordingly. Ratios of normalized peak intensities were fitted for η and δ values as described for pSRII above.

*Maltose binding protein* – ^15^N-TROSY sampled ^15^N T_1_ relaxation experiments^51^ were collected at 750 MHz with relaxation delays of 0.000, 0.240 (2x), 0.400, 0.560, 0.800 (2x), 1.040, 1.360, 1.680 (2x), and 2.000 s. ^15^N-TROSY-sampled T_1ρ_ relaxation experiments^51^ were also collected at 750 MHz with relaxation delays of 0.0010, 0.0058 (2x), 0.0106, 0.0163, 0.0228 (2x), 0.0300, 0.0380, 0.0460, and 0.0550 s. Decay curves for resolved peaks in the ^15^N-TROSY spectrum were analyzed identically. The global reorientation time for MBP in H_2_O buffer were determined as for pSRII above (16.2 ± 0.7 ns). Single- and triple-quantum ^1^H-^13^C coherence transfer experiments were also collected at 750 MHz at a series of delay times: 0.001, 0.004, 0.008 (2x), 0.012, 0.016, 0.020, 0.024 (2x), 0.028, 0.032, and 0.034 s. More scans (48/FID) were acquired for the triple quantum experiment than for the single-quantum experiment (16/FID) and the raw peak intensities normalized accordingly. Ratios of normalized peak intensities were fitted for η and δ values as described for pSRII above. Methyl side chain order parameters were calculated as above for pSRII; however, because the experiments were collected in D_2_O buffer, the global reorientation time was multiplied by 1.223 as to account for the known viscosity difference in the solvents.

**Molecular dynamics simulations**

pSRII (PDB:1H68^7^) was protonated with VMD^56^ and parameters/topology for the covalently attached retinal cofactor (Lys205) were used as deposited in the NAMD^57^ repository. The protein, retinal, and its structural water molecules were inserted into an 8.0 by 8.0 nm bilayer composed of 1-palmitoyl-2-oleoyl-sn-glycero-3-phosphocholine (POPC) generated in VMD to maintain nonpolar and aromatic molecules in the bilayer.^56^ The protein-containing bilayer was then hydrated with an additional 8 Å of TIP3P water^58^ on either side along with the removal of any lipid molecules overlapping the protein. All other aspects of the simulation were as previously described^59,60^ for 1000 steps of minimization, 200 ps of equilibration, and 100 total ns of simulation at 50 °C (as in experiment).

Two molecular dynamics simulations^61^ of OmpW (PDB:2MHL^10^) (one in a bilayer and one in a detergent micelle) were built using the CHARMM-GUI server.^62^ The bilayer system was built using 1,2-dimyristoyl-sn-glycero-3-phosphocholine (DMPC). DMPC is well characterized by simulations and is known to match the hydrophobic thickness of the native outer membrane.^63,64^ The micelle system was built using the SB3-12 detergent.

Simulation systems were neutralized using 150 mM NaCl and under NPT conditions (40°C and 1 atm, as in experiment). Simulations were run using NAMD^57^ employing the CHARMM36 forcefield^65^ with 2.0 femtosecond (fs) time steps. The standard equilibration protocol provided by CHARMM-GUI was followed. Following equilibration, the first 50 ns of simulation were used to equilibrate the bilayer or micelle atoms. Simulation temperature was controlled using the Nose-Hoover thermostat, with a damping coefficient of 1.0 ps^-1^, and the bath was not coupled to hydrogen atoms. Pressure was controlled using the Langevin piston with a constant pressure piston of 50 fs and decay time equal to 25 fs. Long range electrostatics were evaluated using the Particle Mesh Ewald summation.^66^ Both short range electrostatics and Leonard Jones interactions were cutoff at 12 Å, with a smooth switching function applied beginning at 10 Å. Non-bonded pairs were determined using a distance cutoff of 16 Å, and the list was updated every 20 femtoseconds. Long range and electrostatics interactions were evaluated every step. The final trajectories were run for 300 ns.

Backbone O^2^ and side chain methyl O^2^_axis_ values were calculated as previously described^60^ using the long-time limit approximation. All 100 ns of pSRII simulation time was included in the calculation (~5x 𝜏_c_) while the 250 ns of simulation for OmpW was split into two separate 125 ns (also ~5x 𝜏_c_) trajectories for analysis.

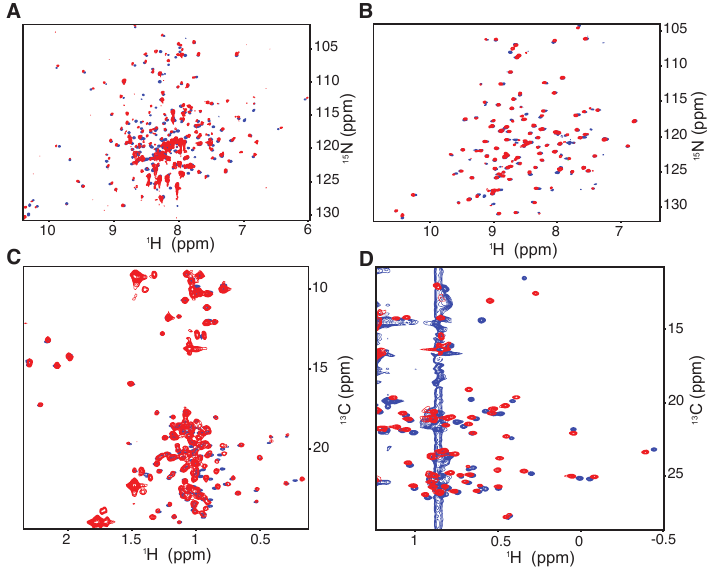

Fig. S1. Comparison of NMR spectra of pSRII & OmpW in micelles and bicelles. Micelle-incorporated spectra are displayed in blue as compared to bicelle-incorporated data in red. pSRII was incorporated into d_26_-c7-DHPC micelles and DMPC/c6-DHPC bicelles at a q-value of 1.1 while OmpW was incorporated into SB3-12 detergent micelles and DMPC/SB3-12 bicelles, also with a q-value of 1.1. ^15^N-TROSY spectra of pSRII (50 °C) and OmpW (40 °C) are shown in panels (A) & (B), respectively, while methyl ^13^C-^1^H HMQC spectra of pSRII and OmpW are shown in panels (C) & (D), respectively. Streaks from detergent molecules are more pronounced in (D) for OmpW due to the lack of availability of deuterated SB3-12.

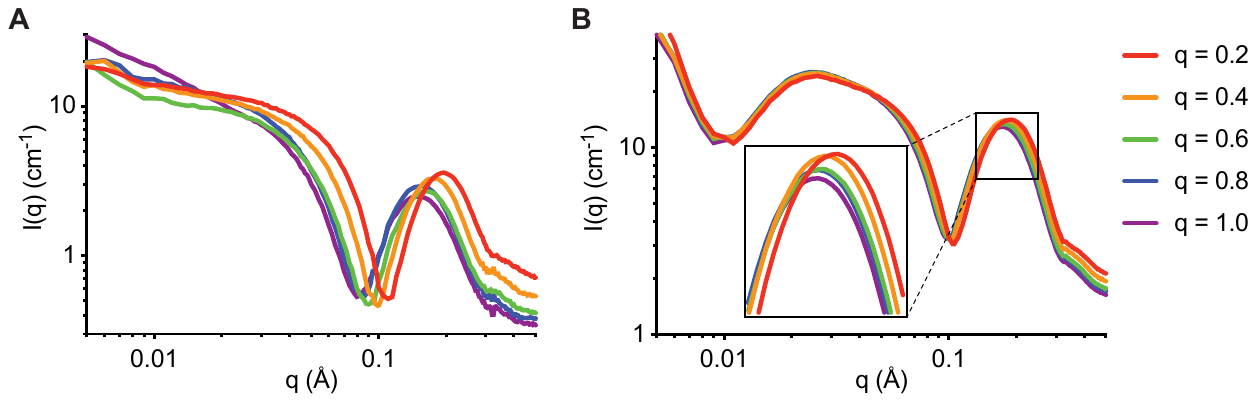

**Fig. S2. Small-angle X-ray scattering characterization of bicelle-incorporated membrane proteins at varying q-values. (A)** PSRII samples scattered as a function of q ratio (ratio of DMPC to c6-DHPC). The q = 1.0 bicelle sample was measured via SAS (purple). Subsequent samples with lower q-ratios were assembled via addition of c6-DHPC and measured with the following q ratios: 0.8 (blue), 0.6 (green), 0.4 (orange), 0.2 (red). The change in the prominent peak feature, as observed by Q_max_ (Q value at maximum value) is highlighted with arrows. (**B**) Similarly, OmpW was measured at a series of q-ratios with the same color scheme as for pSRII. Zooming in on the major peak feature in the OmpW SAXS highlights the similar relationship between bicelle q-ratio and peak feature behavior.

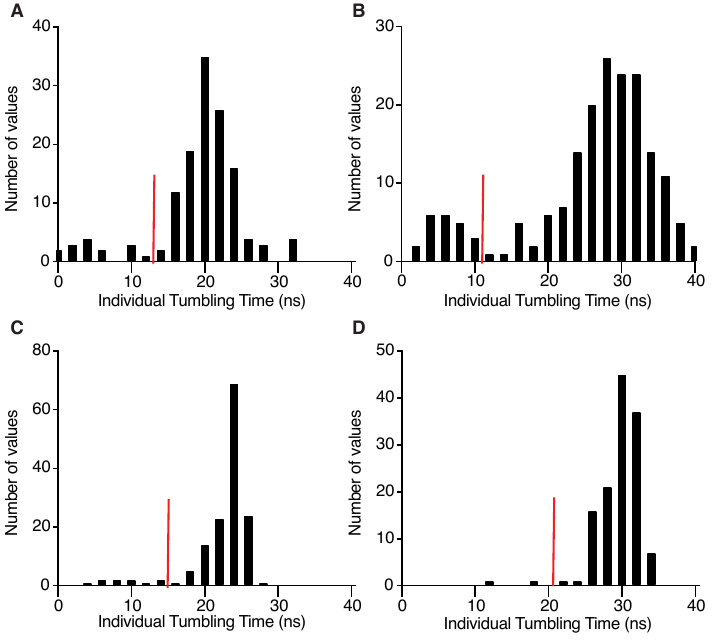

Fig. S3. Distribution of effective tumbling times for pSRII and OmpW. The ^15^N amide T_1_/T_2_ ratio at each residue was used to determine local effective tumbling times for pSRII incorporated into detergent micelles (A) and bicelles (B). Shown here is the distribution obtained using data collected at 750 MHz (^1^H) (micelles, A) and at 800 MHz (^1^H) (bicelles, B) for the same sample used subsequently for cross-correlated relaxation experiments. The red lines demarcate outliers discarded before averaging. OmpW incorporated into micelles (C) and q=1.1 bicelles (D) were analyzed comparably and the local effective tumbling times for individual residues are plotted above. Shown here are the distributions obtained using data collected at 600 MHz (^1^H) (micelles, C) and at 800 MHz (^1^H) (bicelles, D). pSRII has an apparently broader distribution in local tumbling times due to incomplete deuteration and lower local precision.

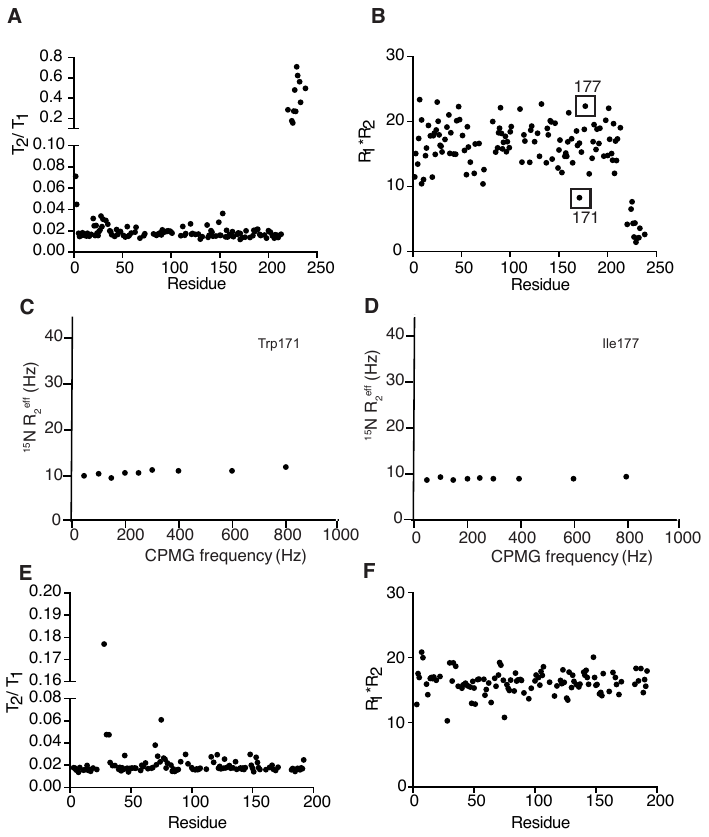

**Fig. S4. The helical and sheet backbones of pSRII and OmpW are largely dynamically silent**. (**A**) T_1_/T_2_ ratios and (**B**) R_1_*R_2_ products averages for experiments collected on pSRII at both 600 and 750 MHz. Example CPMG dispersion curves are shown in (**C**) and (**D**) for example outlier residues in the R_1_*R_2_ product (**B**) demonstrating a lack of intermediate-timescale exchange motions in pSRII. (**E**) T_1_/T_2_ ratios and (**F**) R_1_*R_2_ products for experiments collected on OmpW at 600 MHz demonstrate a comparable lack of fast and intermediate timescale motions.

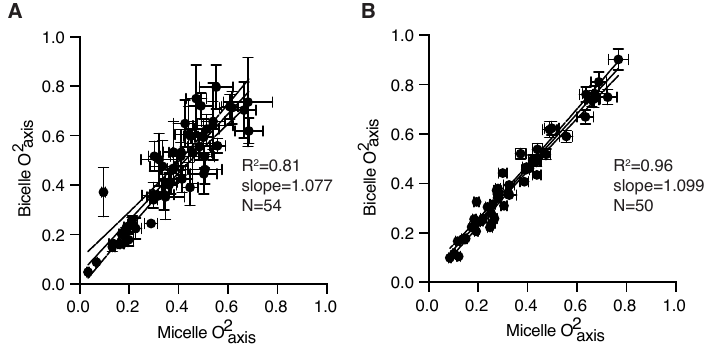

**Fig. S5. Correlation of O^2^_axis_ values for pSRII & OmpW in detergent micelles & lipid bilayer bicelles.** Both pSRII (**A**) and OmpW (**B**) display clear, linear correlations of experimentally determined methyl O^2^_axis_ values when incorporated into detergent micelles and lipid bilayers. The O^2^_axis_ value for all methyl groups that are easily mapped in both lipid conditions and that display reasonable error values (<0.15) are plotted above with horizontal and vertical error bars representing the error for that particular resonance as determined by Monte Carlo fitting for the micelle and bicelle-incorporated proteins, respectively. Linear fit statistics are shown for each. Errors are much higher in bicelle-incorporated pSRII due to incomplete levels of deuteration.

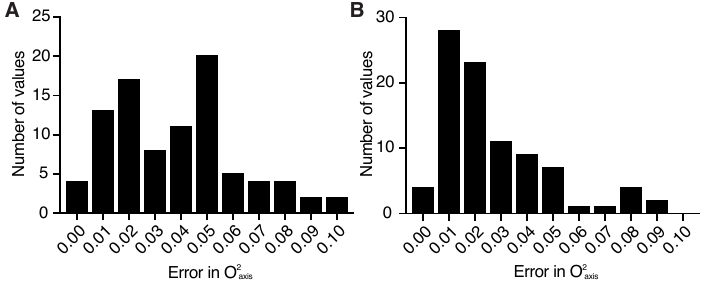

Fig. S6. Distribution in errors in O^2^_axis_ values estimated by Monte Carlo simulations for pSRII at 50 °C (A) and OmpW at 40 °C (B) incorporating experimental fitting error and error in global tumbling time. All O^2^_axis_ values used for this analysis have an error less than 0.10 and have an average error of 0.038 (pSRII) and 0.019 (OmpW).

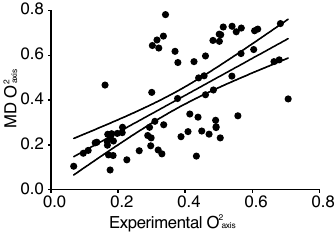

Fig. S7. Comparison with O^2^_axis_ values derived from molecular dynamics simulations of pSRII. Linear correlation of molecular dynamics derived O^2^_axis_ values with those obtained experimentally using cross-correlated relaxation NMR relaxation. The agreement is relatively poor (R^2^ = 0.41, r.m.s.d = 0.16, slope = 0.83, y-intercept = 0.09) and is at the lower end of reliability of similar simulations for soluble proteins.^60^ The 95% confidence windows for the line of best fit are shown surrounding the main fit.

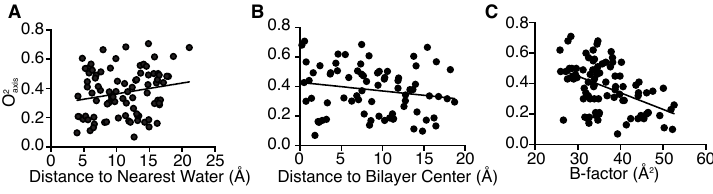

Fig. S8. Potential structural correlates to observed methyl dynamics. (A) Correlation of methyl group motion with distance of the methyl carbon to the nearest water oxygen at the molecular surface (R^2^ = 0.04). (B) Correlation of methyl group motion with distance of the methyl carbon to the putative bilayer center. (C) Correlation of methyl group motion with crystallographic B-factor (R^2^ = 0.214). The crystal structure^7^ (PDB code 1H68) was used. The analysis for depth of burial of each probe was carried out with the TravelDepth program.^67^ Shown are the best-fitted lines to each data set.

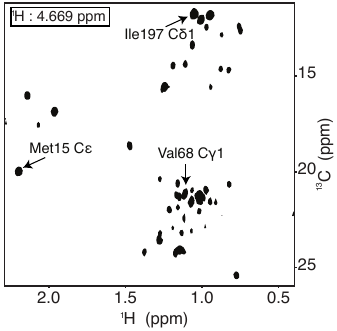

Fig. S9. Protein-water contacts. Two-dimensional plane at the water resonance of a three-dimensional methyl ^13^C-resolved ^1^H-^1^H NOESY spectrum. Most peaks arise from solvent-exposed methyl groups, while the boxed residues of interest represent deeply buried methyl groups within close proximity to both structural water networks discussed in the text.

Table S1. Methyl relaxation data and determined O^2^_axis_ parameters for pSRII in DHPC micelles at 50 °C at 750 MHz (^1^H) (deposited to the BMRB under accession number 27465).

| **Atom** | **δ^[a]^** | **η^[a]^** | **η error^[b]^** | **Reduced χ^2[c,d]^** | **O^2^_axis_ ^[d]^** | **O^2^_axis_ error ^[e]^** |
| --- | --- | --- | --- | --- | --- | --- |
| Met1 Cε | 0.30 | 2.65 | 0.37 | 1.34 | 0.034 | 0.005 |
| Val2 Cγ1 | -4.23 | 7.64 | 0.58 | 0.46 | 0.097 | 0.008 |
| Val2 Cγ2 | -16.17 | 8.81 | 0.95 | 0.30 | 0.112 | 0.013 |
| Leu4 Cδ1 | -25.19 | 16.31 | 3.37 | 1.11 | 0.206 | 0.044 |
| Leu4 Cδ2 | -13.39 | 16.08 | 0.63 | 0.13 | 0.203 | 0.011 |
| Leu7 Cδ2 | -36.18 | 23.89 | 1.68 | 0.44 | 0.302 | 0.024 |
| Leu10 Cδ1 | -10.72 | 13.30 | 0.90 | 1.04 | 0.168 | 0.013 |
| Ile13 Cδ1 | -16.07 | 23.75 | 0.82 | 2.31 | 0.301 | 0.016 |
| Leu16 Cδ1 | -8.90 | 13.33 | 0.93 | 1.14 | 0.169 | 0.014 |
| Leu20 Cδ1 | -20.82 | 22.80 | 1.83 | 1.60 | 0.288 | 0.025 |
| Val38 Cγ1 | -17.53 | 30.65 | 1.11 | 0.70 | 0.388 | 0.022 |
| Val38 Cγ2 | -32.28 | 32.71 | 3.09 | 1.42 | 0.414 | 0.042 |
| Leu40 Cδ2 | -43.24 | 34.64 | 7.59 | 1.59 | 0.438 | 0.099 |
| Ile46 Cδ1 | -38.72 | 36.41 | 3.52 | 2.22 | 0.461 | 0.050 |
| Val49 Cγ2 | -54.83 | 54.05 | 3.98 | 0.77 | 0.684 | 0.057 |
| Val53 Cγ1 | -31.64 | 38.79 | 3.79 | 2.75 | 0.491 | 0.051 |
| Val53 Cγ2 | -31.81 | 37.25 | 3.13 | 0.76 | 0.471 | 0.042 |
| Leu56 Cδ1 | -17.34 | 12.38 | 1.53 | 0.98 | 0.157 | 0.021 |
| Leu56 Cδ2 | -44.44 | 39.98 | 5.22 | 1.91 | 0.506 | 0.069 |
| Val58 Cγ1 | -26.55 | 42.66 | 3.20 | 1.29 | 0.540 | 0.048 |
| Val58 Cγ2 | -35.83 | 47.96 | 1.53 | 0.80 | 0.607 | 0.033 |
| Val61 Cγ1 | -32.57 | 32.44 | 3.57 | 0.73 | 0.411 | 0.050 |
| Val63 Cγ1 | -27.93 | 35.26 | 2.01 | 0.25 | 0.446 | 0.031 |
| Val63 Cγ2 | -61.96 | 42.48 | 3.29 | 2.00 | 0.538 | 0.047 |
| Val68 Cγ1 | -36.02 | 47.73 | 3.85 | 3.58 | 0.604 | 0.054 |
| Ile74 Cδ1 | -16.88 | 24.48 | 0.73 | 1.08 | 0.310 | 0.016 |
| Ile77 Cδ1 | -43.39 | 44.02 | 1.95 | 0.96 | 0.557 | 0.033 |
| Leu78 Cδ1 | -49.62 | 44.75 | 4.42 | 0.89 | 0.566 | 0.060 |
| Leu78 Cδ2 | -32.17 | 33.68 | 3.46 | 0.82 | 0.426 | 0.048 |
| Val84 Cγ1 | -41.46 | 39.65 | 3.91 | 1.01 | 0.502 | 0.054 |
| Val84 Cγ2 | -51.89 | 38.15 | 3.14 | 0.83 | 0.483 | 0.046 |
| Leu87 Cδ1 | -37.55 | 39.63 | 2.32 | 0.53 | 0.502 | 0.036 |
| Leu89 Cδ1 | -27.41 | 28.52 | 2.14 | 0.36 | 0.361 | 0.031 |
| Leu89 Cδ2 | -31.87 | 29.99 | 3.00 | 0.39 | 0.380 | 0.041 |
| Leu90 Cδ1 | -26.21 | 23.22 | 2.50 | 0.66 | 0.294 | 0.035 |
| Leu90 Cδ2 | -28.50 | 26.16 | 3.79 | 0.94 | 0.331 | 0.050 |
| Leu93 Cδ1 | -35.44 | 26.49 | 3.94 | 0.81 | 0.335 | 0.050 |
| Leu93 Cδ2 | -33.92 | 24.98 | 4.25 | 0.96 | 0.316 | 0.058 |
| Ile100 Cδ1 | -21.06 | 29.84 | 1.23 | 1.61 | 0.378 | 0.023 |
| Val101 Cγ1 | -18.16 | 14.05 | 1.26 | 0.60 | 0.178 | 0.018 |
| Val101 Cγ2 | -24.17 | 14.31 | 2.03 | 1.22 | 0.181 | 0.025 |
| Ile102 Cδ1 | -17.06 | 13.40 | 0.56 | 0.65 | 0.170 | 0.010 |
| Val107 Cγ1 | -51.98 | 55.85 | 3.30 | 0.68 | 0.707 | 0.051 |
| Met109 Cε | -82.07 | 53.77 | 7.27 | 0.55 | 0.680 | 0.099 |
| Met117 Cε | -4.03 | 13.57 | 0.21 | 0.16 | 0.172 | 0.008 |
| Val118 Cγ1 | -40.94 | 41.54 | 4.45 | 1.30 | 0.526 | 0.062 |
| Ile121 Cδ1 | -4.51 | 14.99 | 0.49 | 2.18 | 0.190 | 0.010 |
| Met129 Cε | -2.75 | 5.38 | 0.19 | 0.47 | 0.068 | 0.004 |
| Ile135 Cδ1 | -21.84 | 25.25 | 1.46 | 1.72 | 0.320 | 0.023 |
| Leu137 Cδ1 | -38.96 | 29.25 | 4.00 | 0.61 | 0.370 | 0.054 |
| Leu137 Cδ2 | -37.35 | 25.39 | 5.75 | 1.99 | 0.321 | 0.072 |
| Val138 Cγ1 | -55.28 | 43.65 | 4.85 | 0.41 | 0.552 | 0.068 |
| Val138 Cγ2 | -45.71 | 48.69 | 3.45 | 0.78 | 0.616 | 0.050 |
| Leu141 Cδ1 | -27.69 | 23.51 | 3.37 | 0.89 | 0.298 | 0.046 |
| Leu141 Cδ2 | -9.02 | 14.74 | 0.46 | 0.58 | 0.187 | 0.010 |
| Val142 Cγ2 | -25.83 | 27.00 | 2.84 | 0.73 | 0.342 | 0.040 |
| Met145 Cε | -39.95 | 29.90 | 3.95 | 1.32 | 0.378 | 0.051 |
| Ile156 Cδ1 | -16.59 | 16.69 | 0.62 | 0.74 | 0.211 | 0.012 |
| Val161 Cγ1 | -36.85 | 44.82 | 4.38 | 0.92 | 0.567 | 0.061 |
| Val161 Cγ2 | -46.86 | 40.60 | 3.61 | 0.46 | 0.514 | 0.049 |
| Val163 Cδ2 | -50.51 | 36.08 | 6.45 | 0.64 | 0.457 | 0.078 |
| Leu166 Cδ2 | -34.01 | 16.80 | 2.78 | 0.77 | 0.213 | 0.039 |
| Leu166 Cδ1 | -19.04 | 26.57 | 1.89 | 1.42 | 0.336 | 0.028 |
| Val168 Cγ1 | -30.94 | 39.89 | 2.84 | 1.20 | 0.505 | 0.040 |
| Leu170 Cδ2 | -7.90 | 12.75 | 0.99 | 0.39 | 0.161 | 0.015 |
| Ile173 Cδ1 | -13.47 | 23.61 | 0.80 | 2.70 | 0.299 | 0.016 |
| Ile177 Cδ1 | -21.01 | 17.22 | 1.14 | 0.70 | 0.218 | 0.017 |
| Leu179 Cδ1 | -9.93 | 16.88 | 1.43 | 0.90 | 0.214 | 0.020 |
| Leu179 Cδ2 | -14.36 | 15.63 | 1.44 | 0.97 | 0.198 | 0.020 |
| Val185 Cγ1 | -21.51 | 29.13 | 1.34 | 0.34 | 0.369 | 0.025 |
| Val185 Cγ2 | -47.29 | 27.46 | 5.75 | 0.89 | 0.347 | 0.075 |
| Leu187 Cδ1 | -7.26 | 14.68 | 0.58 | 0.37 | 0.186 | 0.011 |
| Leu187 Cδ2 | -6.09 | 10.43 | 0.71 | 0.90 | 0.132 | 0.011 |
| Leu188 Cδ2 | -45.87 | 42.60 | 6.46 | 1.76 | 0.539 | 0.081 |
| Val192 Cγ2 | -62.96 | 52.54 | 3.04 | 0.15 | 0.665 | 0.048 |
| Leu196 Cδ1 | -33.12 | 32.27 | 2.88 | 2.88 | 0.408 | 0.041 |
| Leu196 Cδ2 | -34.89 | 34.26 | 6.13 | 2.80 | 0.434 | 0.080 |
| Ile197 Cδ1 | -16.90 | 10.85 | 1.35 | 0.83 | 0.137 | 0.018 |
| Val198 Cγ1 | -37.19 | 36.38 | 3.30 | 0.79 | 0.460 | 0.045 |
| Leu200 Cδ1 | -72.41 | 38.26 | 6.89 | 0.56 | 0.484 | 0.092 |
| Leu202 Cδ2 | -35.26 | 34.79 | 4.11 | 1.40 | 0.440 | 0.054 |
| Val203 Cγ1 | -33.56 | 38.91 | 5.14 | 1.53 | 0.492 | 0.066 |
| Val203 Cγ2 | -43.20 | 38.91 | 2.90 | 1.12 | 0.492 | 0.045 |
| Val206 Cγ1 | -41.09 | 48.94 | 6.92 | 2.50 | 0.619 | 0.094 |
| Ile211 Cδ1 | -33.43 | 35.51 | 1.73 | 1.36 | 0.449 | 0.029 |
| Leu213 Cδ1 | -21.88 | 20.83 | 1.30 | 0.53 | 0.264 | 0.020 |
| Leu213 Cδ2 | -17.78 | 17.91 | 1.70 | 1.35 | 0.227 | 0.024 |
| Leu219 Cδ2 | -10.78 | 16.50 | 1.71 | 1.60 | 0.209 | 0.024 |
| Val230 Cγ1 | -12.60 | 1.80 | 0.20 | 0.55 | 0.023 | 0.003 |
| Val237 Cγ1 | -12.02 | 1.74 | 0.25 | 0.92 | 0.022 | 0.003 |

^[a]^ η and δ were fitted using Eq. 1. ^[b]^	Errors in η were estimated from the covariance in fitting to Eq. 1 and incorporated the reduced χ^2^ values. ^[c]^	Reduced χ^2^ for the fit with Eq. 1. ^[d]^	O^2^_axis_ values were determined from Eq. 2 using the fitted η values and the calculated tumbling molecular reorientation time (21.9 ns for the data in this Table). ^[e]^	Errors in final O^2^_axis_ were determined by Monte Carlo simulations incorporating errors in fitted η as well as the error in the determined reorientation time.

Table S2. Methyl relaxation data and determined O^2^_axis_ parameters for OmpW in SB3-12 micelles at 40 °C at 600 MHz (^1^H).

| **Peak Number** | **δ^[a]^** | **η^[a]^** | **η error^[b]^** | **Reduced χ^2[c,d]^** | **O^2^_axis_ ^[d]^** | **O^2^_axis_ error ^[e]^** |
| --- | --- | --- | --- | --- | --- | --- |
| a1 | -13.17 | 20.74 | 0.26 | 0.46 | 0.238 | 0.008 |
| a2 | -12.24 | 22.02 | 0.45 | 1.11 | 0.253 | 0.010 |
| a3 | -15.31 | 16.84 | 0.59 | 2.34 | 0.193 | 0.010 |
| a4 | -9.36 | 22.50 | 0.41 | 1.18 | 0.258 | 0.010 |
| a5 | -13.14 | 24.25 | 0.44 | 0.94 | 0.279 | 0.010 |
| a6 | -11.12 | 24.42 | 0.61 | 0.97 | 0.281 | 0.011 |
| a7 | -13.04 | 24.18 | 0.39 | 0.58 | 0.278 | 0.010 |
| a8 | -14.26 | 19.20 | 0.67 | 0.95 | 0.221 | 0.011 |
| a9 | -3.94 | 7.45 | 0.34 | 1.79 | 0.086 | 0.005 |
| a10 | -5.42 | 10.55 | 0.47 | 1.32 | 0.121 | 0.007 |
| a11 | -33.89 | 42.63 | 2.14 | 1.64 | 0.490 | 0.029 |
| a12 | -8.91 | 8.90 | 0.15 | 0.49 | 0.102 | 0.004 |
| a13 | -35.02 | 48.87 | 1.57 | 1.11 | 0.561 | 0.026 |
| a14 | -42.51 | 58.48 | 3.44 | 2.21 | 0.672 | 0.049 |
| a15 | -24.27 | 38.34 | 1.35 | 1.10 | 0.440 | 0.021 |
| a16 | -28.84 | 38.04 | 1.02 | 0.45 | 0.437 | 0.019 |
| a17 | -49.49 | 66.86 | 2.98 | 1.12 | 0.768 | 0.041 |
| a19 | -6.21 | 13.42 | 0.40 | 0.89 | 0.154 | 0.007 |
| a20 | -21.90 | 34.47 | 0.90 | 0.70 | 0.396 | 0.017 |
| a21 | -31.34 | 38.11 | 0.96 | 0.67 | 0.438 | 0.018 |
| a23 | -31.54 | 63.07 | 2.48 | 0.63 | 0.725 | 0.037 |
| a25 | -32.07 | 57.52 | 1.51 | 0.55 | 0.661 | 0.028 |
| a26 | -34.61 | 59.96 | 1.86 | 1.00 | 0.689 | 0.032 |
| a27 | -12.79 | 23.18 | 0.34 | 0.24 | 0.266 | 0.010 |
| a28 | -14.17 | 21.60 | 0.40 | 0.22 | 0.248 | 0.009 |
| a29 | -23.33 | 43.35 | 2.10 | 0.95 | 0.498 | 0.029 |
| a31 | -35.61 | 55.43 | 1.45 | 0.66 | 0.637 | 0.027 |
| a33 | -15.49 | 32.39 | 1.59 | 1.46 | 0.372 | 0.021 |
| a34 | -45.31 | 48.46 | 1.68 | 0.48 | 0.557 | 0.026 |
| a36 | -10.47 | 26.07 | 1.22 | 0.68 | 0.299 | 0.018 |
| a38 | -14.31 | 41.41 | 0.41 | 0.44 | 0.476 | 0.017 |
| a40 | -23.64 | 23.87 | 0.43 | 1.08 | 0.274 | 0.011 |
| a41 | -8.37 | 13.01 | 0.20 | 1.02 | 0.149 | 0.005 |
| a43 | -19.70 | 35.51 | 0.63 | 0.91 | 0.408 | 0.015 |
| a44 | -9.94 | 15.12 | 0.54 | 4.47 | 0.174 | 0.009 |
| a45 | -29.46 | 55.21 | 1.97 | 1.96 | 0.634 | 0.033 |
| a47 | -6.10 | 17.68 | 0.27 | 0.78 | 0.203 | 0.007 |
| a48 | -9.67 | 28.29 | 1.69 | 18.03 | 0.325 | 0.023 |
| a49 | -14.11 | 37.66 | 0.48 | 1.87 | 0.433 | 0.015 |
| a50 | -9.41 | 28.09 | 0.23 | 0.24 | 0.323 | 0.011 |
| a51 | -10.64 | 26.52 | 0.51 | 1.14 | 0.305 | 0.012 |
| a52 | -17.58 | 34.18 | 0.96 | 1.37 | 0.393 | 0.016 |
| a59 | -14.70 | 33.70 | 1.06 | 0.45 | 0.387 | 0.017 |
| a60 | -9.58 | 12.19 | 0.48 | 3.67 | 0.140 | 0.007 |
| a61 | -8.34 | 28.40 | 2.25 | 1.05 | 0.326 | 0.030 |
| a62 | -20.63 | 36.69 | 0.65 | 0.70 | 0.421 | 0.017 |
| a100 | -31.11 | 58.20 | 1.34 | 2.87 | 0.669 | 0.027 |
| a101 | -7.45 | 18.56 | 0.16 | 0.56 | 0.213 | 0.007 |
| a103 | -11.40 | 22.20 | 0.88 | 5.70 | 0.255 | 0.014 |
| a104 | -8.59 | 16.71 | 0.39 | 0.40 | 0.192 | 0.007 |
| a105 | -14.58 | 10.19 | 0.71 | 1.08 | 0.117 | 0.009 |
| a107 | -9.75 | 15.98 | 0.39 | 0.74 | 0.184 | 0.008 |

^[a]^ η and δ were fitted using Eq. 1. ^[b]^	Errors in η were estimated from the covariance in fitting to Eq. 1 and incorporated the reduced χ^2^ values. ^[c]^	Reduced χ^2^ for the fit with Eq. 1. ^[d]^	O^2^_axis_ values were determined from Eq. 2 using the fitted η values and the calculated tumbling molecular reorientation time (24.0 ns for the data in this Table). ^[e]^	Errors in final O^2^_axis_ were determined by Monte Carlo simulations incorporating errors in fitted η as well as the error in the determined reorientation time.

Table S3: Methyl relaxation data and determined O^2^_axis_ parameters for pSRII in DMPC/DHPC bicelles at 50 °C at 800 MHz (^1^H)

| **Atom** | **δ^[a]^** | **η^[a]^** | **η error^[b]^** | **Reduced χ^2[c,d]^** | **O^2^_axis_ ^[d]^** | **O^2^_axis_ error ^[e]^** |
| --- | --- | --- | --- | --- | --- | --- |
| Met1 Cε | -44.38 | 5.07 | 1.42 | 0.33 | 0.049 | 0.013 |
| Val2 Cγ1 | -76.63 | 38.85 | 10.81 | 0.86 | 0.372 | 0.099 |
| Leu7 Cδ2 | -57.96 | 54.01 | 6.26 | 0.27 | 0.517 | 0.060 |
| Ile13 Cδ1 | -26.22 | 37.82 | 0.98 | 0.29 | 0.362 | 0.011 |
| Met15 Cε | -141.7 | 78.50 | 15.63 | 0.20 | 0.752 | 0.149 |
| Leu20 Cδ1 | -8.59 | 25.59 | 1.52 | 0.17 | 0.245 | 0.015 |
| Val38 Cγ2 | -39.66 | 44.39 | 5.75 | 0.72 | 0.425 | 0.054 |
| Ile43 Cδ1 | -95.25 | 61.97 | 4.22 | 0.05 | 0.594 | 0.044 |
| Ile46 Cδ1 | -62.34 | 64.72 | 5.35 | 0.69 | 0.620 | 0.052 |
| Val49 Cγ2 | -39.40 | 75.31 | 5.01 | 0.24 | 0.721 | 0.052 |
| Val53 Cγ2 | -53.87 | 78.34 | 14.65 | 0.75 | 0.751 | 0.137 |
| Leu56 Cδ2 | -25.08 | 48.43 | 5.95 | 0.48 | 0.464 | 0.060 |
| Val58 Cγ1 | -86.65 | 68.71 | 15.92 | 0.34 | 0.658 | 0.156 |
| Val58 Cγ2 | -51.52 | 74.91 | 5.27 | 0.44 | 0.718 | 0.049 |
| Val61 Cγ1 | -48.88 | 55.66 | 9.67 | 0.51 | 0.533 | 0.091 |
| Val63 Cγ1 | -16.96 | 40.86 | 7.40 | 0.32 | 0.391 | 0.073 |
| Ile77 Cδ1 | -43.94 | 58.40 | 2.95 | 0.46 | 0.559 | 0.030 |
| Leu78 Cδ2 | -51.07 | 67.83 | 9.99 | 0.70 | 0.650 | 0.095 |
| Val84 Cγ1 | -32.82 | 46.60 | 8.52 | 1.38 | 0.446 | 0.081 |
| Val84 Cγ2 | -59.51 | 58.07 | 5.58 | 0.28 | 0.556 | 0.054 |
| Leu87 Cδ1 | -38.49 | 62.09 | 7.71 | 0.51 | 0.595 | 0.072 |
| Leu89 Cδ1 | -36.36 | 41.85 | 5.10 | 0.17 | 0.401 | 0.049 |
| Leu89 Cδ2 | -66.20 | 55.71 | 13.46 | 0.69 | 0.534 | 0.125 |
| Leu93 Cδ1 | -55.40 | 49.70 | 10.59 | 0.23 | 0.476 | 0.105 |
| Ile100 Cδ1 | -33.04 | 48.81 | 2.11 | 0.54 | 0.468 | 0.020 |
| Ile102 Cδ1 | -20.78 | 21.15 | 1.39 | 0.34 | 0.203 | 0.013 |
| Met109 Cε | -136.0 | 76.82 | 18.42 | 0.72 | 0.736 | 0.180 |
| Met117 Cε | -2.89 | 17.03 | 1.13 | 0.60 | 0.163 | 0.011 |
| Ile121 Cδ1 | -10.60 | 24.17 | 0.50 | 0.49 | 0.232 | 0.006 |
| Met129 Cε | -9.60 | 9.25 | 0.68 | 1.06 | 0.089 | 0.007 |
| Ile135 Cδ1 | -28.05 | 37.34 | 1.38 | 0.25 | 0.358 | 0.014 |
| Leu137 Cδ2 | -55.71 | 52.58 | 10.66 | 0.29 | 0.504 | 0.106 |
| Val138 Cγ1 | -76.86 | 83.22 | 9.85 | 0.22 | 0.797 | 0.089 |
| Val138 Cγ2 | -57.22 | 74.50 | 6.61 | 0.51 | 0.714 | 0.066 |
| Leu141 Cδ1 | -30.20 | 37.54 | 5.32 | 0.25 | 0.360 | 0.052 |
| Leu141 Cδ2 | -11.80 | 21.82 | 1.98 | 0.34 | 0.209 | 0.020 |
| Val142 Cγ2 | -37.16 | 37.81 | 6.31 | 0.51 | 0.362 | 0.062 |
| Met145 Cε | -64.12 | 48.40 | 5.82 | 0.36 | 0.464 | 0.058 |
| Ile156 Cδ1 | -26.15 | 25.77 | 2.25 | 0.82 | 0.247 | 0.022 |
| Val161 Cγ2 | -61.09 | 65.47 | 5.89 | 0.11 | 0.627 | 0.058 |
| Val163 Cγ2 | -50.69 | 55.60 | 12.59 | 0.21 | 0.533 | 0.126 |
| Val168 Cγ1 | -32.77 | 53.80 | 4.52 | 0.42 | 0.515 | 0.045 |
| Ile173 Cδ1 | -24.92 | 35.62 | 0.95 | 0.50 | 0.341 | 0.011 |
| Leu179 Cδ1 | -15.40 | 25.38 | 3.59 | 0.41 | 0.243 | 0.033 |
| Leu179 Cδ2 | -0.11 | 18.73 | 2.58 | 0.31 | 0.179 | 0.025 |
| Val185 Cγ1 | -29.39 | 43.03 | 4.82 | 0.71 | 0.412 | 0.047 |
| Val185 Cγ2 | -31.34 | 36.82 | 9.71 | 0.46 | 0.353 | 0.090 |
| Leu187 Cδ1 | -0.23 | 17.99 | 1.47 | 0.21 | 0.172 | 0.015 |
| Leu187 Cδ2 | -14.90 | 15.76 | 2.00 | 0.32 | 0.151 | 0.019 |
| Leu188 Cδ2 | -73.68 | 66.96 | 10.03 | 0.38 | 0.641 | 0.096 |
| Val192 Cγ2 | -105.2 | 73.49 | 11.80 | 0.21 | 0.704 | 0.113 |
| Leu196 Cδ1 | -57.96 | 54.01 | 6.26 | 0.27 | 0.517 | 0.056 |
| Leu196 Cδ2 | -31.50 | 37.59 | 3.92 | 0.10 | 0.360 | 0.037 |
| Ile197 Cδ1 | -23.15 | 17.06 | 2.00 | 0.19 | 0.163 | 0.019 |
| Leu202 Cδ2 | -52.44 | 63.23 | 11.58 | 0.36 | 0.606 | 0.113 |
| Ile211 Cδ1 | -59.71 | 62.48 | 2.82 | 0.24 | 0.599 | 0.029 |
| Leu213 Cδ2 | -39.78 | 23.49 | 4.61 | 0.61 | 0.225 | 0.042 |
| Leu219 Cδ2 | -26.24 | 32.23 | 4.42 | 0.25 | 0.309 | 0.044 |

^[a]^ η and δ were fitted using Eq. 1. ^[b]^	Errors in η were estimated from the covariance in fitting to Eq. 1 and incorporated the reduced χ^2^ values. ^[c]^	Reduced χ^2^ for the fit with Eq. 1. ^[d]^	O^2^_axis_ values were determined from Eq. 2 using the fitted η values and the calculated tumbling molecular reorientation time (28.9 ns for the data in this Table). ^[e]^	Errors in final O^2^_axis_ were determined by Monte Carlo simulations incorporating errors in fitted η as well as the error in the determined reorientation time.

Table S4: Methyl relaxation data and determined O^2^_axis_ parameters for OmpW in DMPC/DHPC bicelles at 40 °C at 800 MHz (^1^H)

| **Peak Number** | **δ^[a]^** | **η^[a]^** | **η error^[b]^** | **Reduced χ^2[c,d]^** | **O^2^_axis_ ^[d]^** | **O^2^_axis_ error ^[e]^** |
| --- | --- | --- | --- | --- | --- | --- |
| a1 | -12.55 | 33.89 | 0.54 | 10.86 | 0.305 | 0.011 |
| a2 | -12.48 | 31.05 | 0.54 | 9.47 | 0.279 | 0.011 |
| a3 | -15.16 | 36.17 | 0.54 | 6.14 | 0.325 | 0.012 |
| a4 | -10.34 | 35.14 | 0.53 | 9.86 | 0.316 | 0.012 |
| a5 | -15.36 | 37.96 | 0.75 | 12.62 | 0.341 | 0.013 |
| a6 | -18.16 | 42.22 | 0.82 | 4.37 | 0.379 | 0.015 |
| a7 | -14.28 | 39.71 | 0.68 | 6.26 | 0.357 | 0.014 |
| a8 | -14.42 | 28.81 | 0.44 | 3.28 | 0.259 | 0.010 |
| a9 | -7.00 | 10.96 | 0.09 | 0.99 | 0.098 | 0.003 |
| a10 | -8.17 | 11.62 | 0.20 | 6.03 | 0.104 | 0.004 |
| a11 | -43.61 | 68.78 | 2.80 | 2.87 | 0.618 | 0.032 |
| a12 | -7.16 | 12.86 | 0.26 | 2.77 | 0.116 | 0.005 |
| a14 | -44.89 | 85.03 | 2.13 | 2.48 | 0.764 | 0.031 |
| a15 | -35.23 | 59.86 | 1.29 | 1.71 | 0.538 | 0.023 |
| a16 | -32.78 | 54.39 | 1.34 | 2.52 | 0.489 | 0.021 |
| a17 | -52.39 | 100.38 | 3.23 | 2.89 | 0.902 | 0.043 |
| a18 | -58.34 | 87.24 | 3.83 | 3.72 | 0.784 | 0.042 |
| a20 | -25.36 | 51.86 | 1.04 | 2.36 | 0.466 | 0.018 |
| a21 | -24.60 | 48.40 | 1.04 | 3.71 | 0.435 | 0.018 |
| a22 | -47.16 | 89.70 | 2.18 | 0.62 | 0.806 | 0.033 |
| a23 | -39.82 | 83.44 | 1.94 | 0.82 | 0.750 | 0.030 |
| a24 | -48.46 | 80.21 | 2.78 | 2.25 | 0.721 | 0.033 |
| a25 | -37.03 | 82.22 | 1.42 | 0.66 | 0.739 | 0.028 |
| a26 | -50.03 | 90.28 | 2.71 | 3.44 | 0.811 | 0.039 |
| a27 | -16.93 | 28.73 | 0.38 | 1.45 | 0.258 | 0.009 |
| a28 | -16.44 | 24.71 | 0.23 | 0.39 | 0.222 | 0.008 |
| a29 | -16.13 | 69.40 | 1.81 | 0.70 | 0.624 | 0.028 |
| a30 | -33.11 | 54.33 | 1.54 | 2.14 | 0.488 | 0.021 |
| a31 | -40.27 | 84.61 | 1.90 | 3.60 | 0.760 | 0.031 |
| a33 | -7.49 | 57.90 | 1.19 | 0.81 | 0.520 | 0.022 |
| a34 | -42.67 | 65.66 | 1.69 | 1.47 | 0.590 | 0.024 |
| a36 | -16.30 | 49.19 | 0.80 | 0.19 | 0.442 | 0.016 |
| a38 | -9.58 | 57.60 | 1.29 | 21.16 | 0.518 | 0.021 |
| a40 | -15.86 | 41.36 | 0.31 | 1.94 | 0.372 | 0.013 |
| a41 | -3.58 | 19.61 | 0.26 | 2.13 | 0.176 | 0.007 |
| a43 | -21.01 | 51.35 | 1.22 | 13.17 | 0.461 | 0.019 |
| a44 | -8.67 | 24.92 | 0.55 | 14.70 | 0.224 | 0.009 |
| a45 | -35.80 | 74.53 | 2.15 | 7.04 | 0.670 | 0.029 |
| a46 | -7.81 | 24.40 | 0.46 | 4.73 | 0.219 | 0.008 |
| a47 | -6.64 | 27.31 | 0.44 | 10.15 | 0.245 | 0.009 |
| a48 | -10.18 | 43.83 | 0.82 | 14.74 | 0.394 | 0.015 |
| a49 | -8.51 | 54.69 | 0.81 | 15.25 | 0.491 | 0.019 |
| a50 | -7.63 | 40.23 | 0.61 | 9.09 | 0.362 | 0.014 |
| a51 | -6.89 | 34.42 | 0.43 | 3.89 | 0.309 | 0.011 |
| a52 | -22.76 | 50.66 | 0.97 | 4.47 | 0.455 | 0.018 |
| a59 | -13.99 | 45.32 | 1.14 | 5.00 | 0.407 | 0.018 |
| a60 | -3.38 | 19.48 | 0.30 | 2.78 | 0.175 | 0.007 |
| a61 | -19.27 | 39.24 | 1.06 | 2.56 | 0.353 | 0.016 |
| a62 | -19.65 | 54.99 | 1.27 | 8.41 | 0.494 | 0.021 |
| a100 | -27.17 | 82.57 | 2.79 | 22.35 | 0.742 | 0.035 |
| a101 | -7.26 | 27.63 | 0.38 | 12.14 | 0.248 | 0.009 |
| a103 | -16.74 | 26.61 | 0.39 | 2.28 | 0.239 | 0.009 |
| a104 | -9.29 | 22.98 | 0.29 | 2.40 | 0.207 | 0.008 |
| a105 | -16.22 | 18.47 | 0.45 | 0.72 | 0.166 | 0.007 |
| a107 | -5.60 | 28.44 | 0.58 | 1.90 | 0.256 | 0.011 |

^[a]^ η and δ were fitted using Eq. 1. ^[b]^	Errors in η were estimated from the covariance in fitting to Eq. 1 and incorporated the reduced χ^2^ values. ^[c]^	Reduced χ^2^ for the fit with Eq. 1. ^[d]^	O^2^_axis_ values were determined from Eq. 2 using the fitted η values and the calculated tumbling molecular reorientation time (29.9 ns for the data in this Table). ^[e]^	Errors in final O^2^_axis_ were determined by Monte Carlo simulations incorporating errors in fitted η as well as the error in the determined reorientation time.

Table S5: Methyl relaxation data and determined O^2^_axis_ parameters for maltose binding protein at 37 °C at 750 MHz (^1^H)

| **Atom** | **δ^[a]^** | **η^[a]^** | **η error^[b]^** | **Reduced χ^2[c,d]^** | **O^2^_axis_ ^[d]^** | **O^2^_axis_ error ^[e]^** |
| --- | --- | --- | --- | --- | --- | --- |
| 1 | -4.50 | 11.20 | 0.20 | 3.15 | 0.157 | 0.006 |
| 2 | -19.45 | 39.99 | 0.37 | 1.26 | 0.559 | 0.021 |
| 3 | -6.72 | 28.60 | 0.29 | 3.42 | 0.400 | 0.014 |
| 4 | -2.88 | 14.76 | 0.21 | 3.74 | 0.206 | 0.008 |
| 5 | -15.24 | 49.15 | 1.15 | 11.70 | 0.687 | 0.029 |
| 6 | -3.28 | 29.42 | 1.28 | 81.35 | 0.411 | 0.023 |
| 7 | -6.66 | 19.22 | 0.35 | 7.83 | 0.269 | 0.011 |
| 8 | -17.10 | 39.19 | 0.64 | 4.84 | 0.548 | 0.022 |
| 9 | -27.32 | 56.69 | 1.23 | 5.12 | 0.792 | 0.032 |
| 10 | -12.87 | 36.24 | 0.54 | 6.15 | 0.506 | 0.019 |
| 11 | -20.03 | 54.82 | 1.54 | 13.83 | 0.766 | 0.035 |
| 12 | -8.97 | 24.23 | 0.18 | 1.39 | 0.339 | 0.012 |
| 13 | -20.39 | 53.59 | 1.23 | 3.45 | 0.749 | 0.033 |
| 14 | -6.43 | 22.18 | 0.22 | 1.62 | 0.310 | 0.011 |
| 15 | -19.43 | 41.16 | 0.70 | 4.45 | 0.575 | 0.023 |
| 16 | -12.80 | 36.63 | 0.56 | 4.57 | 0.512 | 0.020 |
| 17 | -30.88 | 56.23 | 1.59 | 15.87 | 0.786 | 0.033 |
| 18 | -20.20 | 43.96 | 0.62 | 5.51 | 0.614 | 0.023 |
| 19 | -4.91 | 37.97 | 1.14 | 26.54 | 0.531 | 0.024 |
| 20 | -32.21 | 54.06 | 1.69 | 1.08 | 0.755 | 0.036 |
| 21 | -21.12 | 54.35 | 1.50 | 3.54 | 0.759 | 0.034 |
| 22 | -4.50 | 37.70 | 1.42 | 22.82 | 0.527 | 0.027 |
| 23 | -22.05 | 60.89 | 1.10 | 1.92 | 0.851 | 0.033 |
| 24 | 0.33 | 9.60 | 0.19 | 2.37 | 0.134 | 0.005 |
| 25 | -22.17 | 57.61 | 1.31 | 3.75 | 0.805 | 0.034 |
| 26 | -4.31 | 37.61 | 1.66 | 39.47 | 0.526 | 0.030 |
| 27 | -18.30 | 58.99 | 2.09 | 5.42 | 0.824 | 0.042 |
| 28 | -13.70 | 61.10 | 1.29 | 3.03 | 0.854 | 0.033 |
| 29 | -15.55 | 61.01 | 2.09 | 8.40 | 0.853 | 0.042 |
| 30 | -16.35 | 60.72 | 2.00 | 7.84 | 0.848 | 0.041 |
| 31 | -15.78 | 53.34 | 0.78 | 1.59 | 0.745 | 0.028 |
| 32 | -7.11 | 34.30 | 0.75 | 3.79 | 0.479 | 0.019 |
| 33 | -18.10 | 54.69 | 1.48 | 5.26 | 0.764 | 0.034 |
| 34 | -11.13 | 52.43 | 1.05 | 3.09 | 0.733 | 0.030 |
| 35 | -22.57 | 64.43 | 1.39 | 1.92 | 0.900 | 0.037 |
| 36 | -4.90 | 41.43 | 1.23 | 5.35 | 0.579 | 0.025 |
| 37 | -16.07 | 60.28 | 1.93 | 7.00 | 0.842 | 0.041 |
| 38 | -11.44 | 58.24 | 2.15 | 8.01 | 0.814 | 0.043 |
| 39 | -13.05 | 51.16 | 1.66 | 5.72 | 0.715 | 0.033 |
| 40 | -15.02 | 58.86 | 1.48 | 3.21 | 0.823 | 0.034 |
| 41 | -42.10 | 66.17 | 2.70 | 5.12 | 0.925 | 0.049 |
| 42 | -8.65 | 28.60 | 0.55 | 3.10 | 0.400 | 0.016 |
| 44 | -13.16 | 62.73 | 2.32 | 8.13 | 0.877 | 0.045 |
| 46 | -6.44 | 38.25 | 1.12 | 14.09 | 0.535 | 0.024 |
| 47 | -18.73 | 41.88 | 1.07 | 2.94 | 0.585 | 0.025 |
| 48 | -19.09 | 44.48 | 0.73 | 1.94 | 0.622 | 0.024 |
| 49 | -5.93 | 34.08 | 0.89 | 10.22 | 0.476 | 0.021 |
| 50 | -15.42 | 62.02 | 2.17 | 12.04 | 0.867 | 0.045 |
| 51 | -16.90 | 50.33 | 1.28 | 4.00 | 0.703 | 0.030 |
| 52 | -30.88 | 59.84 | 1.98 | 2.77 | 0.836 | 0.042 |
| 53 | -8.59 | 44.00 | 1.03 | 6.70 | 0.615 | 0.027 |
| 54 | -26.25 | 56.35 | 1.43 | 2.37 | 0.788 | 0.035 |
| 55 | -24.16 | 47.76 | 1.24 | 1.84 | 0.667 | 0.029 |
| 56 | -9.80 | 30.08 | 0.49 | 2.36 | 0.420 | 0.017 |
| 57 | -22.85 | 43.34 | 0.90 | 1.57 | 0.606 | 0.024 |
| 58 | -29.34 | 53.46 | 1.81 | 3.75 | 0.747 | 0.036 |
| 59 | -13.28 | 50.72 | 1.86 | 8.71 | 0.709 | 0.036 |
| 60 | -13.29 | 57.41 | 1.43 | 5.98 | 0.802 | 0.036 |

^[a]^ η and δ were fitted using Eq. 1. ^[b]^	Errors in η were estimated from the covariance in fitting to Eq. 1 and incorporated the reduced χ^2^ values. ^[c]^	Reduced χ^2^ for the fit with Eq. 1. ^[d]^	O^2^_axis_ values were determined from Eq. 2 using the fitted η values and the calculated tumbling molecular reorientation time (19.8 ns for the data in this Table, calculated from 16.2 ns in H_2_O multiplied by 1.223 to account for the viscosity difference in the D_2_O buffer used here). ^[e]^	Errors in final O^2^_axis_ were determined by Monte Carlo simulations incorporating errors in fitted η as well as the error in the determined reorientation time.

Table S6. Soluble protein average methyl O^2^_axis_ values for isoleucine, leucine, valine and methionine residues.^[a]^

**Abbreviation	Protein^[ref]^	ILVM <O^2^_axis_>	Temperature (°C)**

Ca^2+^-CaM	Ca^2+^ – saturated calmodulin (CaM)^4^	0.427	35

smMLCK-CaM	CaM – smMLCK(p) complex)^27^	0.478	50

nNOS-CaM	CaM – nNOS(p) complex^4^	0.499	35

ubq	ubiquitin^68^	0.505	45

smMLCK-CaM	CaM – smMLCK(p) complex^4^	0.532	35

CaMKKa-CaM	CaM – CaMKKa(p) complex^4^	0.544	35

PhoB-DNA	DNA binding domain PhoB – DNA^69^	0.587	37

MBP	Maltose binding protein^[b]^	0.611	37

MSG	malate synthase G^70^	0.656	37

Flavodoxin	Bacterial flavodoxin C55A^71^	0.668	35

HEWL	hen egg white lysozyme^72^	0.677	35

^[a]^ Source data for Fig. 2

^[b]^ This work (Table S5)

**Additional References**

44.	Burgess, N. K., Dao, T. P., Stanley, A. M. & Fleming, K. G. B-Barrel Proteins That Reside in the Escherichia coli Outer Membrane in Vivo Demonstrate Varied Folding Behavior. *J. Biol. Chem.* **283**, 26748–26758 (2008).

45.	Goto, N. K., Gardner, K. H., Mueller, G. A., Willis, R. C. & Kay, L. E. A robust and cost-effective method for the production of Val, Leu, Ile (??1) methyl-protonated 15N-, 13C-, 2H-labeled proteins. *J. Biomol. NMR* **13**, 369–374 (1999).

46.	Moon, C. P., Zaccai, N. R., Fleming, P. J., Gessmann, D. & Fleming, K. G. Membrane protein thermodynamic stability may serve as the energy sink for sorting in the periplasm. *Proc Natl Acad Sci U S A* **110**, 4285–4290 (2013).

47.	Gardner, K. H. & Kay, L. E. Production and Incorporation of 15N, 13C, 2H (1H-d1 Methyl) Isoleucine into Proteins for Multidimensional NMR Studies. *J. Am. Chem. Soc.* **119**, 7599–7600 (1997).

48.	Difabio, J. *et al.* The life science x-ray scattering beamline at NSLS-II. *AIP Conf. Proc.* **1741**, (2016).

49.	Hyberts, S. G., Milbradt, A. G., Wagner, A. B., Arthanari, H. & Wagner, G. Application of iterative soft thresholding for fast reconstruction of NMR data non-uniformly sampled with multidimensional poisson gap scheduling. *J. Biomol. NMR* **52**, 315–327 (2012).

50.	Delaglio, F. *et al.* NMRPipe : A multidimensional spectral processing system based on UNIX pipes. *J. Biomol. NMR* **6**, 277–293 (1995).

51.	Lakomek, N.-A., Ying, J. & Bax, A. Measurement of 15N relaxation rates in perdeuterated proteins by TROSY-based methods. *J. Biomol. NMR* **53**, 209–221 (2012).

52.	Cavanagh, J., Fairbrother, W. J., Palmer, A. G., Rance, M. & Skelton, N. J. *Protein NMR Spectroscopy: Principles and Practice*. (Elsevier, 2007).

53.	Loria, J. P., Rance, M. & Palmer, A. G. A TROSY CPMG sequence for characterizing chemical exchange in large proteins. *J. Biomol. NMR* **15**, 151–155 (1999).

54.	Korzhnev, D. M., Kloiber, K., Kanelis, V., Tugarinov, V. & Kay, L. E. Probing Slow Dynamics in High Molecular Weight Proteins by Methyl-TROSY NMR Spectroscopy : Application to a 723-Residue Enzyme. *J Am Chem Soc* **126**, 3964–3973 (2004).

55.	Fu, Y. *et al.* Coupled motion in proteins revealed by pressure perturbation SI. *J. Am. Chem. Soc.* **134**, 8543–8550 (2012).

56.	Humphrey, W., Dalke, A. & Schulten, K. VMD : Visual Molecular Dynamics. *J. Mol. Graph.* **14**, 33–38 (1996).

57.	Phillips, J. C. *et al.* Scalable molecular dynamics with NAMD. *J. Comput. Chem.* **26**, 1781–1802 (2005).

58.	Jorgensen, W. L., Chandrasekhar, J., Madura, J. D., Impey, R. W. & Klein, M. L. Comparison of simple potential functions for simulating liquid water. *J. Chem. Phys.* **79**, 926 (1983).

59.	Kasinath, V., Sharp, K. A. & Wand, A. J. Microscopic insights into the NMR relaxation-based protein conformational entropy meter. *J. Am. Chem. Soc.* **135**, 15092–100 (2013).

60.	O’Brien, E. S., Wand, A. J. & Sharp, K. A. On the ability of molecular dynamics force fields to recapitulate NMR derived protein side chain order parameters. *Protein Sci.* **25**, 1156–60 (2016).

61.	Towns, J. *et al.* XSEDE: Accelerating Scientific Discovery. *Comput. Sci. Eng.* 62–74 (2014).

62.	Lee, J. *et al.* CHARMM-GUI Input Generator for NAMD, GROMACS, AMBER, OpenMM, and CHARMM/OpenMM Simulations Using the CHARMM36 Additive Force Field. *J. Chem. Theory Comput.* **12**, 405–413 (2016).

63.	Klauda, J. B. *et al.* E. coli Outer Membrane and Interactions with OmpLA. *Biophys. J.* **106**, 2493–2502 (2014).

64.	Kucerka, N. *et al.* Structure of Fully Hydrated Fluid Phase DMPC and DLPC Lipid Bilayers Using X-Ray Scattering from Oriented Multilamellar Arrays and from Unilamellar Vesicles. *Biophys. J.* **88**, 2626–2637 (2005).

65.	Klauda, J. B. *et al.* Update of the CHARMM All-Atom Additive Force Field for Lipids : Validation on Six Lipid Types. *J. Phys. Chem. B* **114**, 7830–7843 (2010).

66.	York, D. M. & Yang, W. A chemical potential equalization method for molecular simulations A chemical potential equalization method for molecular simulations. *J. Chem. Phys.* **104**, 159–172 (1996).

67.	Coleman, R. G. & Sharp, K. A. Travel Depth, a New Shape Descriptor for Macromolecules: Application to Ligand Binding. *J. Mol. Biol.* **362**, 441–458 (2006).

68.	Wand, A. J., Urbauer, J. L., Mcevoy, R. P. & Bieber, R. J. Internal Dynamics of Human Ubiquitin Revealed by 13C-Relaxation Studies of Randomly Fractionally Labeled Protein. *Biochemistry* **35**, 6116–6125 (1996).

69.	Okamura, H., Makino, K. & Nishimura, Y. NMR Dynamics Distinguish Between Hard and Soft Hydrophobic Cores in the DNA-binding Domain of PhoB and Demonstrate Different Roles of the Cores in Binding to DNA. *J. Mol. Biol.* **367**, 1093–1117 (2007).

70.	Tugarinov, V. & Kay, L. E. Quantitative 13 C and 2 H NMR Relaxation Studies of the 723-Residue Enzyme Malate Synthase G Reveal a Dynamic Binding Interface. *Biochemistry* **44**, 15970–15977 (2005).

71.	Liu, W. *et al.* Main Chain and Side Chain Dynamics of Oxidized Flavodoxin from Cyanobacterium anabaena. *Biochemistry* **40**, 14744–14753 (2001).

72.	Moorman, V. R., Valentine, K. G. & Wand, A. J. The dynamical response of hen egg white lysozyme to the binding of a carbohydrate ligand. *Protein Sci.* **21**, 1066–1073 (2012).

73.	Frauenfelder, H., Sligar, S. G. & Wolynes, P. G. The energy landscapes and motions of proteins. *Science* **254**, 1598–1603 (1991).
